# Radial-glia-to-astrocyte trans-differentiation and astrocyte transcriptional convergence are coordinated by CEH-43/DLX in *C. elegans*

**DOI:** 10.1101/2025.10.17.682973

**Authors:** Simin Liu, Kenneth Bradley, Jinghong J. Tang, Yoon A. Kim, Ana Milosevic, Shai Shaham

## Abstract

Mammalian radial glia can remodel to become astrocytes, which acquire common transcriptional states despite spatially and lineally distinct origins. To uncover molecular programs driving convergent radial-glia-to-astrocyte transformation, we investigated development of *C. elegans* CEPsh astrocytes, which also arise from distinct progenitors without cell division. Using lineage-restricted single-cell RNA sequencing, we delineate a two-phase program for CEPsh astrocyte formation. Transcriptionally disparate nascent CEPsh glia rapidly acquire a common radial-glia-like state, facilitating nerve ring (brain) assembly. Subsequently, convergent CEPsh glia upregulate astrocyte-specific gene expression. Both phases require the *distal-less* transcription factor CEH-43, expressed in CEPsh glia and their progenitors. CEH-43 binds conserved astrocyte-expressed genes, cell-autonomously activating both early and late CEPsh glia-specific gene expression. CEH-43 misexpression is sufficient to induce CEPsh astrocyte reporter expression. We demonstrate that CEH-43 homologs, DLX1/2, are expressed in mouse astrocytes, and comparative transcriptomics reveal additional parallels. Our findings provide a molecular foundation for understanding cell-division-independent radial-glia-to-astrocyte transformation.

## Introduction

Astrocytes are essential for central nervous system (CNS) development and function, supporting neuronal survival(Banker 1980), driving synapse formation(Christopherson et al. 2005; Kucukdereli et al. 2011; Allen et al. 2012; Pfrieger and Barres 1997) and elimination(Chung et al. 2013; Vainchtein et al. 2018), regulating ion and water homeostasis(Olsen et al. 2015; Parpura and Verkhratsky 2012; Kimelberg 2004; Pham et al. 2024), supplying neurons with metabolic substrates and cholesterol(Weber and Barros 2015; Mauch et al. 2001; Goritz et al. 2005; Pfrieger 2003; Meeks and Mennerick 2003; Bélanger et al. 2011), modulating synaptic transmission(Liu et al. 2021), and maintaining blood-brain barrier integrity(Hösli et al. 2022; Manu et al. 2023; Díaz-Castro et al. 2023). Abnormal astrocyte development contributes to neurodevelopmental disorders including Alexander disease(ALEXANDER 1949; Hagemann 2022), Rett syndrome(Ballas et al. 2009; Sun et al. 2023; Postogna et al. 2025), and autism(Allen et al. 2022b; Vakilzadeh and Martinez-Cerdeño 2023), highlighting the importance of understanding how these glia acquire molecular and functional identities.

In vertebrates, radial glia can directly transform into astrocytes without cell division(Voigt 1989; Brunne et al. 2010). Furthermore, lineage tracing reveals that subsets of mouse astrocytes derived from distinct dorsal and ventral radial glia progenitors nonetheless share similar transcriptomes(Bandler et al. 2022). The molecular mechanisms underlying these fundamental aspects of astrocyte development remain unresolved.

*Caenorhabditis elegans* CEPsh glia provide an experimentally tractable system to address both open questions. Nascent CEPsh glia adopt bipolar radial-glia-like morphologies, extending processes that guide pioneer and follower neuron axons into the nerve ring (brain)(Rapti et al. 2017), (Tabata 2015). These cells then undergo a post-mitotic radial-glia-to-astrocyte-like transformation, elaborating ramified processes that ensheathe and penetrate the nerve ring. Mature CEPsh glia cover non-overlapping spatial domains(White et al. 1986), reminiscent of mammalian astrocyte tiling(Ogata and Kosaka 2002). Like vertebrate astrocytes, CEPsh glia promote synaptogenesis(Colón-Ramos et al. 2007), and contact(White et al. 1986) and ensheathe synapses in tripartite-like configurations(Halassa et al. 2007). These glia also express the conserved astrocyte glutamate transporter GLT1/EAAT2(Katz et al. 2019; Purice et al. 2025) whose loss drives repetitive behavior in both *C. elegans* and mice(Katz et al. 2019; Aida et al. 2015). Importantly, while the two dorsal and two ventral CEPsh glia arise from spatially and lineally distinct progenitors, all four cells ultimately adopt similar morphologies and functions and express common reporter genes. While transcriptional regulators affecting CEPsh maturation have been described(Yoshimura et al. 2008), how CEPsh astrocytes arise by cell-division-independent transformation of radial-glia-like precursors and how CEPsh glia acquire common fates has not been investigated.

Here, we apply a lineage-restricted and temporally resolved single-cell RNA sequencing (scRNA-seq) strategy to generate a high-resolution transcriptional map of early CEPsh glia development from their progenitors. We find that CEPsh glia differentiation proceeds in two transcriptionally and functionally distinct phases. In an early convergence phase (phase I), newly generated and transcriptionally distinct dorsal and ventral CEPsh glia adopt a shared embryonic radial-glia-like state within ∼90 minutes, promoting nervous system assembly. In a later maturation phase (phase II), CEPsh glia undergo cell-division–independent transformation into astrocyte-like glia, upregulating genes controlling cell shape, neurotransmission, and behavior. We identify several previously unknown CEPsh glia fate regulators, and demonstrate that, among these, CEH-43, a *distal-less* homeodomain transcription factor, is a key regulator of both phases of CEPsh glia development. *ceh-43* mutants exhibit reduced expression of CEPsh glia gene reporters and show nerve-ring axon guidance defects similar to those observed following embryonic CEPsh glia ablation(Rapti et al. 2017). CEH-43 directly binds key genes required for CEPsh glia development and function, and its mis-expression is sufficient to induce ectopic CEPsh glia reporter expression. Comparative studies uncover transcriptional similarities between CEPsh glia and early mouse and human radial-glia/astrocytes. Notably, we show that the CEH-43 homologs DLX1/2, expressed in vertebrate radial glia(Anderson et al. 1997b, 1997a; Petryniak et al. 2007), are also expressed in mouse astrocytes, suggesting that the molecular programs for astrocyte convergent differentiation and radial-glia-to-astrocyte post-mitotic transformation may be conserved.

## Results

### A developmental transcriptome of early CEPsh glia

scRNA-seq was previously applied to profile transcriptomes of *C. elegans* embryonic cells(Packer et al. 2019), generating datasets that include 127 embryonic CEPsh glia. However, these studies do not resolve CEPsh glia temporal relationships or lineal origins, with transcriptomes of some closely related sister or cousin cells not identified. To generate a high-resolution developmental transcriptome of CEPsh glia, we labeled embryonic sub-lineages that produce CEPsh glia using combinations of fluorescent reporter transgenes (**Fig. 1A,B**; orange labels). Left (L) and right (R) dorsal CEPsh glia lineages were co-labeled with *ceh-32p::H1-Wcherry* and *vab-3::mNeonGreen*, (**Fig. 1A, Supplemental Fig. S1A**), and ventral lineages with *ceh-36p::H1-Wcherry* and *sptf-1::TY1::EGFP* (**Fig. 1B, Supplemental Fig. S1B**). Embryos were synchronized and collected at 240, 270 (ventral only), 330, and 420 min after the first embryonic cell division, spanning the birth time of CEPsh glia at ∼300-min(Sulston et al. 1983). Following embryo dissociation, co-labeled cells were isolated by fluorescence activated cell sorting (FACS) and processed for 10x Chromium scRNA-seq (**Fig. 1C**). Cells from different strains and time points were processed separately to preserve strain and time information (**Supplemental Fig. S2**).

**Figure 1.**
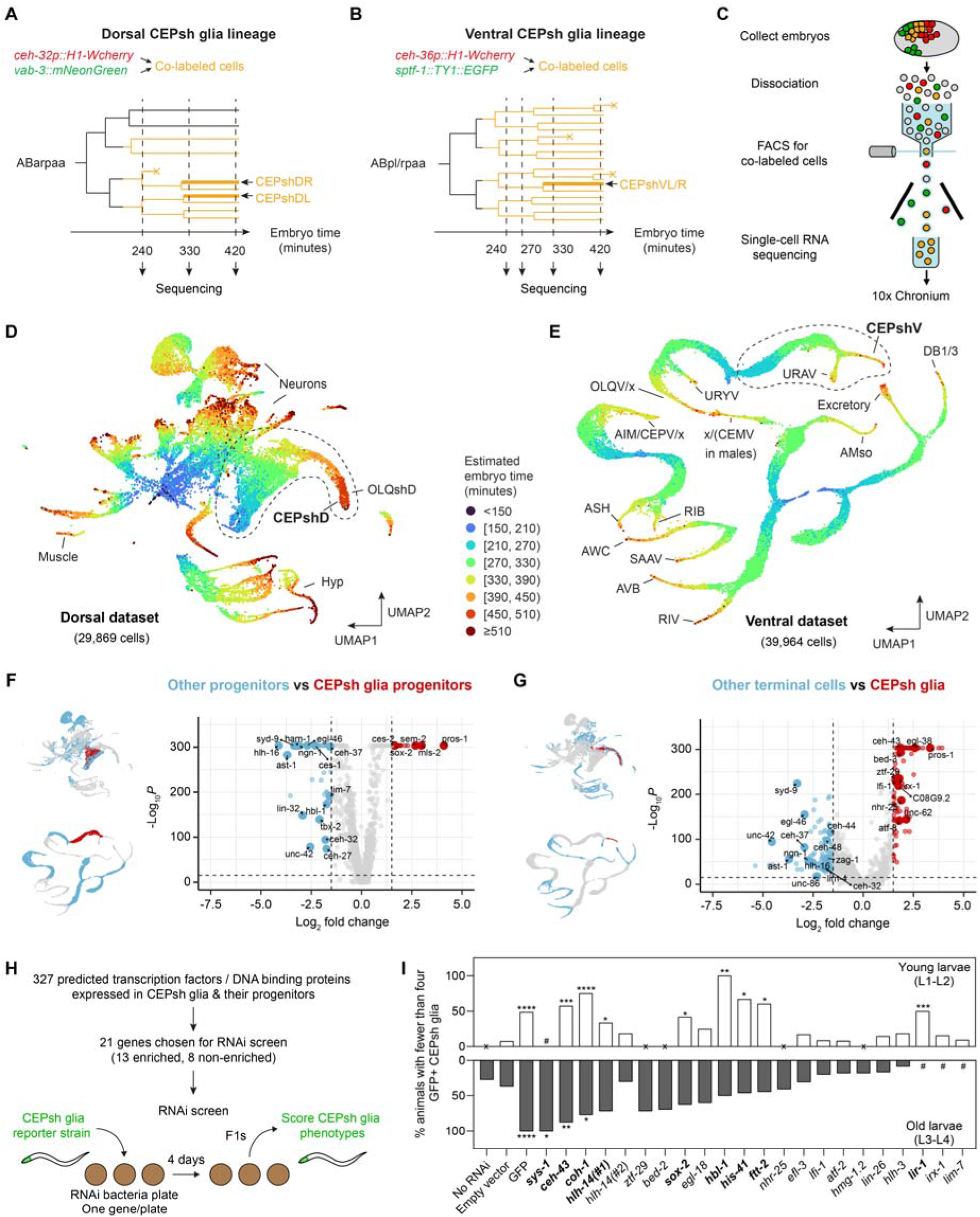
A developmental transcriptome of early CEPsh glia. **A**, Strategy for isolating developing dorsal CEPsh glia. Cells from dorsal CEPsh glia lineages were co-labeled with the two indicated fluorescent reporters. Synchronized embryos were collected at the indicated developmental time points. **B**, Same as (**A**) but for ventral CEPsh glia. **C**, Workflow outlining FACS isolation of co-labeled cells followed by single-cell RNA sequencing. **D**, UMAP projection of 29,869 cells from the dorsal dataset. Each cell is color-coded by the estimated developmental age of the embryo from which the cell was isolated. The sub-lineage containing CEPshD glia and their progenitors is circled. **E**, UMAP projection of the 39,964 cells from the ventral dataset. The sub-lineage containing CEPshV glia and their progenitors is circled. **F**, Differential gene expression analysis comparing CEPsh glia progenitors versus other progenitors. Genes enriched (red) or depleted (blue) in CEPsh glia progenitors are highlighted. Transcription factors are labeled. Dorsal and ventral datasets were combined for analysis. **G**, Same as (**F**), but comparing CEPsh glia with other terminal cell types. **H**, Workflow of the RNAi screen targeting candidate transcription factors and DNA binding proteins. Animals expressing the *hlh-17p::gfp* CEPsh glia reporter were exposed to RNAi, and F1 progeny were scored for GFP expression. **I**, Summary of RNAi screen results. Columns with zero values are marked with “x”. #, phenotype not scored. Fisher’s exact test, \**p* < 0.05, \*\**p* < 0.01, \*\*\**p* < 0.001, \*\*\*\**p* < 0.0001. Raw data, n, and *p*-values are provided in **Supplemental Table S2**.

After quality control (Methods), we retained 29,869 cells from dorsal and 39,964 cells from ventral samples for downstream analysis (**Fig. 1D,E**). To assess the precise embryo age from which individual cells were collected, we compared the transcriptome of each cell with a reference whole-embryo gene expression time course collected every 10 minutes until the onset of muscle movement(Hashimshony et al. 2015). Embryo age was assigned to the time point showing the highest similarity (**Supplemental Fig. S3**). UMAP projections of both dorsal and ventral lineage cells formed continuous trajectories correlating with embryo time (**Fig. 1D,E**, temporal color scale). Consistent with our embryo collection times, 85% of the cells from dorsal samples and 76% of the cells from ventral samples were assigned to 210-450 minutes post-first cell division (**Supplemental Fig. S2C,D**).

Cell types were annotated using published scRNA-seq data(Packer et al. 2019) together with reporter expression patterns (**Supplemental Table S1**). Ventral CEPsh glia (CEPsh**<u>V</u>**) and their progenitors were identified by *pros-1* expression and by their lineage relationship to *cfi-1*-positive URA neurons(Li et al. 2023) (**Supplemental Fig. S4A,B**). To validate our annotations, we imaged the embryonic expression patterns of four reporter transgenes using our custom-built scanning iSPIM microscope(Insley and Shaham 2018; Wu et al. 2011). The observed spatiotemporal patterns largely agree with scRNA-seq, with only minor discrepancies likely reflecting weak reporter expression that falls below the detection threshold of scRNA-seq (**Supplemental Fig. S4C,D**).

The dorsal dataset comprises a broader diversity of cell types. To identify dorsal CEPsh glia (CEPsh**<u>D</u>**) lineages, we first partitioned cells by *pros-1* and *ham-2* expression (**Supplemental Fig. S4E**). *pros-1* was previously shown to be expressed in CEPshD glia and their progenitors ABarpaaaap and ABarpaaapa(Packer et al. 2019; Wallace et al. 2016; Ma et al. 2021), whereas *ham-2* marks the OLQsoD glia progenitor ABarpaaaaa(Packer et al. 2019; Ma et al. 2021) (**Supplemental Fig. S4F,G**). CEPshD glia were then distinguished from their OLQshD glia sister cells based on selective *aqp-7* and *lim-7* expression(Taylor et al. 2021; Ma et al. 2021) (**Supplemental Fig. S4H,I**). We then validated the identities of CEPshD glia and their progenitors using additional published marker genes(Packer et al. 2019) and by confirming that their relative positions on the UMAP projection are consistent with expected lineage architecture (**Fig. 1A,D, Supplemental Fig. S4J-M**).

As independent validation, we identified differentially expressed genes(Hao et al. 2021) between CEPshD/V glia progenitors and other progenitor populations, as well as between CEPshD/V glia and other terminal cell types (**Fig. 1F,G**). We found that cells annotated as CEPsh glia and their progenitors are enriched for known glial regulators (e.g. *pros-1*, *mls-2*)(Yoshimura et al. 2008; Wallace et al. 2016), and depleted for transcription factors that function in neuronal development (e.g. *ngn-1*, *hlh-16*, *ast-1*, *unc-86*)(Nakano et al. 2010; Christensen et al. 2020; Hobert 2016; Flames and Hobert 2009; Bertrand et al. 2011; Sze et al. 2002).

Cells we assigned as CEPsh glia or their progenitors express at least 327 predicted DNA-binding proteins(Weirauch et al. 2014a; Reece-Hoyes et al. 2005a). To test possible roles in CEPsh glia specification, we used RNA interference (RNAi) to knockdown 21 candidates, including 13 enriched in CEPsh glia (average log₂FC > 0, adjusted p-value < 0.05), in RNAi-sensitized *eri-1(mg366)* animals(Kennedy et al. 2004) carrying the mature CEPsh glia reporter *hlh-17p::gfp*(Yoshimura et al. 2008) (**Fig. 1H, Supplemental Table S2**). Knockdown of nine genes significantly reduces *hlh-17p::gfp* expression (**Fig. 1I**), uncovering previously unknown regulators of CEPsh glia fate. Notably, genomic mutations in *lin-26* and *ztf-29* also reduce *hlh-17p::gfp* expression in CEPsh glia despite their relatively weak RNAi phenotypes (**Supplemental Fig. S5**).

Together, these studies delineate a high-resolution transcriptional trajectory for CEPsh glia development, revealing previously undescribed regulators of CEPsh glia specification.

### Nascent CEPsh glia rapidly converge onto a shared radial-glia-like state

How CEPsh glia derived from distinct progenitors ultimately acquire similar morphologies, molecular identities, and functions, remains unknown. To address this question, we examined a joint UMAP embedding of dorsal and ventral CEPsh glia together with their progenitors (**Fig. 2A**). We identified three transcriptionally distinct progenitor types: dorsal left (CEPsh**<u>DL</u>**), dorsal right (CEPsh**<u>DR</u>**), and ventral left/right (CEPsh**<u>V</u>**) progenitors (**Fig. 2A**). CEPshDL and CEPshDR progenitors initially exhibit different transcription profiles, but eventually converge, consistent with their production of transcriptionally indistinguishable CEPshDL/R glia (**Fig. 2A**; lineage inset). Convergence is also evident in transcription factor/DNA-binding protein gene expression (**Fig. 2B**; compare columns *a* vs. *c* and *b* vs. *d*. See **Supplemental Table S3** for the complete gene expression profiles across clusters and **Supplemental Table S13** for a complete list of transcription factors/DNA binding protein genes(Weirauch et al. 2014b; Reece-Hoyes et al. 2005b)) and in global differential gene expression analysis (**Fig. 2C**). By contrast, although no left/right (L/R) differences are detected between CEPshVL and CEPshVR progenitors, these ventral progenitors are markedly different from either CEPshDL/R progenitors (**Fig. 2A,B**; compare column *e* vs. columns *b* or *d*; compare **Fig. 2D,E** vs. **Fig. 2C**). Thus, CEPshD and CEPshV progenitors are transcriptionally distinct.

**Figure 2.**
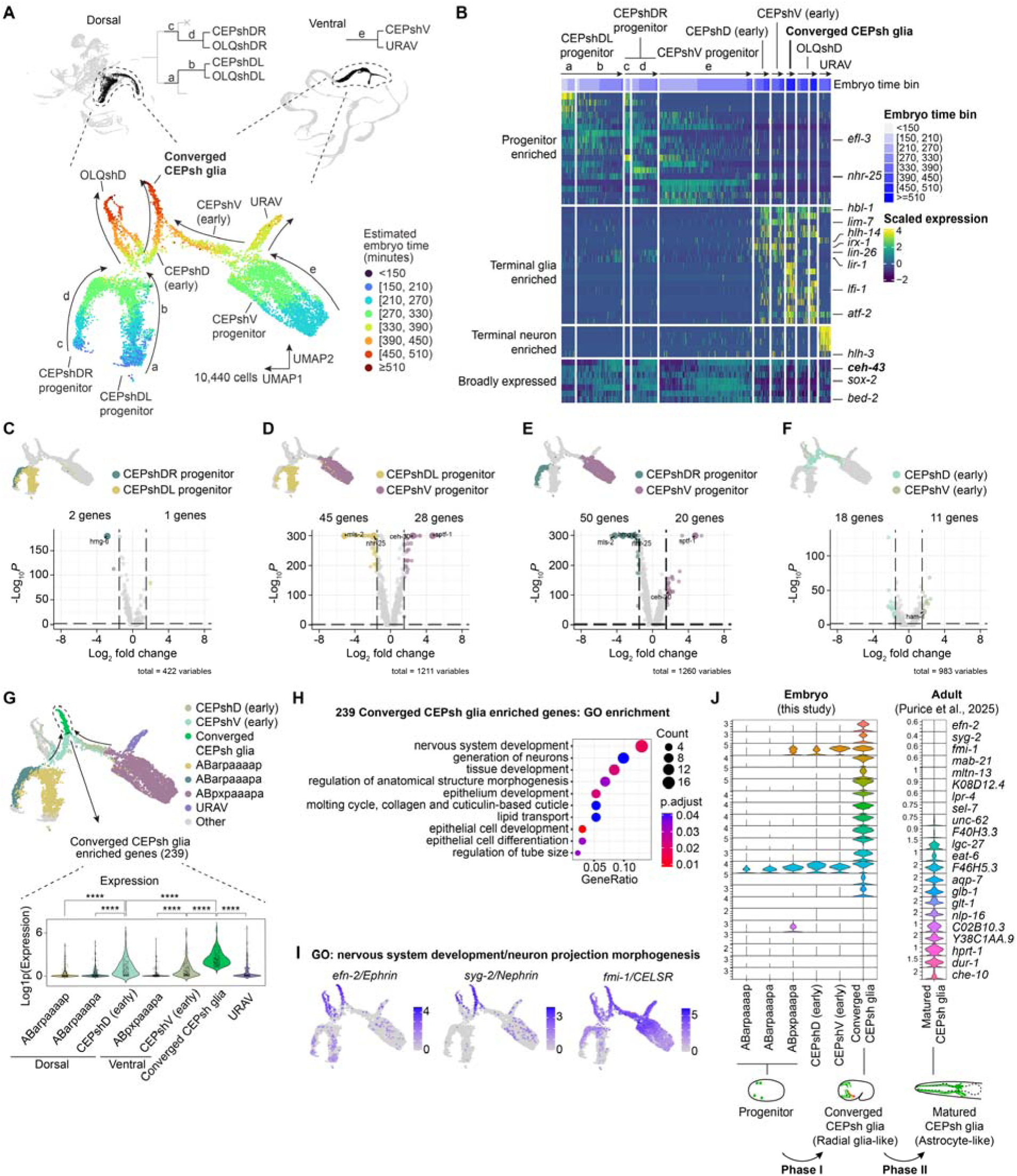
Nascent CEPsh glia converge onto a radial glia-like state, and then undergo a cell division-independent transformation into an astrocyte-like state. **A**, UMAP projection of 10,440 cells from the CEPshD and CEPshV lineages. The lineage relationships are shown in the dendrogram above. Arrows indicate developmental progression over time on the UMAP projection. Lower case letters (a-e) indicate cells whose transcription profiles are shown in panel (**B**). **B**, Heatmap showing expression patterns of transcription factor and DNA binding genes in representative CEPsh glia and their progenitors at different time points. Lower case letters (a-e) correspond to panel (**A**). Expression values were log-transformed, centered, and scaled by the standard deviation of each gene (row). **C-F**, Differential gene expression analysis between the indicated clusters, as illustrated in the corresponding UMAP projections. **G**, Expression dynamics of converged CEPsh glia enriched genes across clusters highlighted in the UMAP projection. For each cluster, the average expression level of each gene is calculated and plotted as an individual dot. \*\*\*\**p* < 0.0001 by Kruskal–Wallis tests with Bonferroni correction for multiple comparisons. **H**, Significantly enriched Gene Ontology (GO) terms among genes upregulated in converged CEPsh glia. **I**, Genes encoding cell membrane proteins implicated in axon guidance and outgrowth (*efn-2*, *fmi-1*) and synapse formation (*syg-*2) are expressed in converged CEPsh glia. **J**, Top, violin plots showing expression dynamics of representative CEPsh glia genes at different developmental stages. Adult CEPsh glia transcriptomic data are from Purice et al. (2025). Bottom, schematic model of CEPsh glia development. In phase I, nascent CEPsh glia converge onto a share state, expressing genes supporting axon guidance and nerve ring assembly (red and yellow indicate pioneer and follower nerve ring neurons), reminiscent of mammalian radial glia. Upon completion of nerve ring formation, phase II begins, marked by the activation of matured astrocyte gene programs, including genes involved in neurotransmitter uptake and broader neuronal activity regulation.

CEPshD glia do not show measurable L/R gene expression differences (**Fig. 2A**; note the single dorsal CEPshD (early) cluster). However, early CEPshD and CEPshV glia are transcriptionally distinct, expressing different transcription factors and other genes (**Fig. 2A,B**,**F**). Remarkably, as early CEPshD and CEPshV glia diverge from their respective sister cells (OLQshD and URAV; **Fig. 2A,B, Supplemental Fig. S6A-C**), they rapidly converge onto a shared transcriptional state within ∼1-1.5 hours (Converged CEPsh glia, **Fig. 2A,B**). The 239 genes enriched in converged CEPsh glia are upregulated in both CEPshD and CEPshV glia (**Fig. 2G**), and expression levels of individual genes are strongly correlated between the two groups (**Supplemental Fig. S6D**), further supporting co-activation of a shared gene program.

Genes enriched in converged CEPsh glia have roles in neurodevelopment and tissue morphogenesis (**Fig. 2H, Supplemental Table S4**). Notably, many, like *efn-2*/Ephrin, *syg-2*/Nephrin, and *fmi-1*/CELSR, encode membrane proteins implicated in axon extension, synaptogenesis, and nerve-ring formation, respectively(Rapti et al. 2017; Shen et al. 2004; Özkan et al. 2014; Sengupta et al. 2021; Tucker et al. 2022; Buyannemekh et al. 2025), processes that we previously showed depend on embryonic CEPsh glia(Rapti et al. 2017) (**Fig. 2I**). Together with their bipolar morphology, this transcriptional signature indicates that converged CEPsh glia acquire a developmental state analogous to vertebrate radial glia, which also extend elongated processes supporting neuronal migration(Rakic 1971; Nadarajah and Parnavelas 2002) and axon pathfinding(Norris and Kalil 1991; Kaur et al. 2020).

### Embryonic radial-glia-like and adult astrocyte-like CEPsh glia transcriptomes are distinct

We compared our embryonic transcriptomes with recently published larval and adult CEPsh glia datasets(Katz et al. 2019; Purice et al. 2025). We found that expression of genes induced in embryonic CEPsh glia and associated with radial-glia-like functions markedly diminishes in adult CEPsh glia (**Fig. 2J**). By contrast, genes characteristic of mature astrocytes, such as the glutamate transporter *glt-1/EAAT2* and the aquaporin *aqp-7*, are upregulated in adult CEPsh glia (**Fig. 2J**). Indeed, 377 genes enriched in L2/L3 larval CEPsh glia(Katz et al. 2019) (log₂FC > 1, adjusted p-value < 0.05) and 323 genes enriched in adult CEPsh glia(Purice et al. 2025) (log₂FC > 0, adjusted p-value < 0.05) (**Supplemental Table S5**) show little or no expression in the embryonic converged CEPsh glia state (**Fig 2J, Supplemental Fig. S6E-G**).

These findings, together with previously described morphological changes between embryonic and adult CEPsh glia, support a two-phase maturation model for CEPsh glia that parallels vertebrate astrocyte development (**Fig. 2J**). In phase I, nascent dorsal and ventral CEPsh glia, generated from distinct progenitors, rapidly converge onto a common radial-glia-like state, upregulating a transcriptional program supporting nerve-ring assembly(Rapti et al. 2017) and presumably required to form and maintain a bipolar morphology. In phase II, these glia undergo a cell-division-independent transformation into adult astrocyte-like glia, marked by extensive morphological complexity and activation of a distinct gene program that, as in mammalian astrocytes, regulates neurotransmission and behavior (**Fig. 2J**).

### CEH-43 promotes convergent CEPsh glia differentiation

Two models can explain how nascent dorsal and ventral CEPsh glia, initially distinct in lineage and transcription, ultimately converge onto a shared gene expression profile. In one model, dorsal- and ventral-specific transcription factors independently activate convergently expressed genes by binding different but functionally equivalent *cis*-regulatory elements. Alternatively, a handful of transcription factors co-expressed in both lineages can bind common regulatory elements to drive a unified transcriptional program. Supporting the latter model, we identified several transcription factors expressed in both CEPshD and CEPshV progenitors and in nascent CEPshD and CEPshV glia (**Fig. 1F,G,2B**). Among these is CEH-43, whose RNAi knockdown reduces expression of the CEPsh glia reporter *hlh-17p::gfp* (**Fig. 1I**).

CEH-43 is a *distal-less* homeobox transcription factor orthologous to mammalian DLX proteins(Aspöck and Bürglin 2001) (**Fig. 3A**). An endogenous *ceh-43::SL2::GFP::H2B* reporter, integrated using CRISPR/Cas9 downstream of the *ceh-43* stop codon, is continuously expressed in all CEPsh glia in the embryo and through adulthood (**Fig. 3B,C**), consistent with *ceh-43* detection in both embryonic and adult CEPsh glia using scRNA-seq(Purice et al. 2025). A *ceh-43(ot406)* missense mutation, predicted to change a single amino acid in the carboxyl terminus of the CEH-43 homeodomain (**Fig. 3A**), strongly reduces *hlh-17p::gfp* expression (**Fig. 3D-F**) and causes defects in CEPsh glia positioning and process morphology (**Supplemental Fig. S7A**). However, *ceh-43(ot406)* mutants maintain normal numbers of head glia scored using the pan-glia reporter *mir-228p::2xNLS::iBlueberry* (**Supplemental Fig. S7B,C**). Thus, CEH-43 is not required for general glial cell production but is essential for CEPsh glia differentiation.

**Figure 3.**
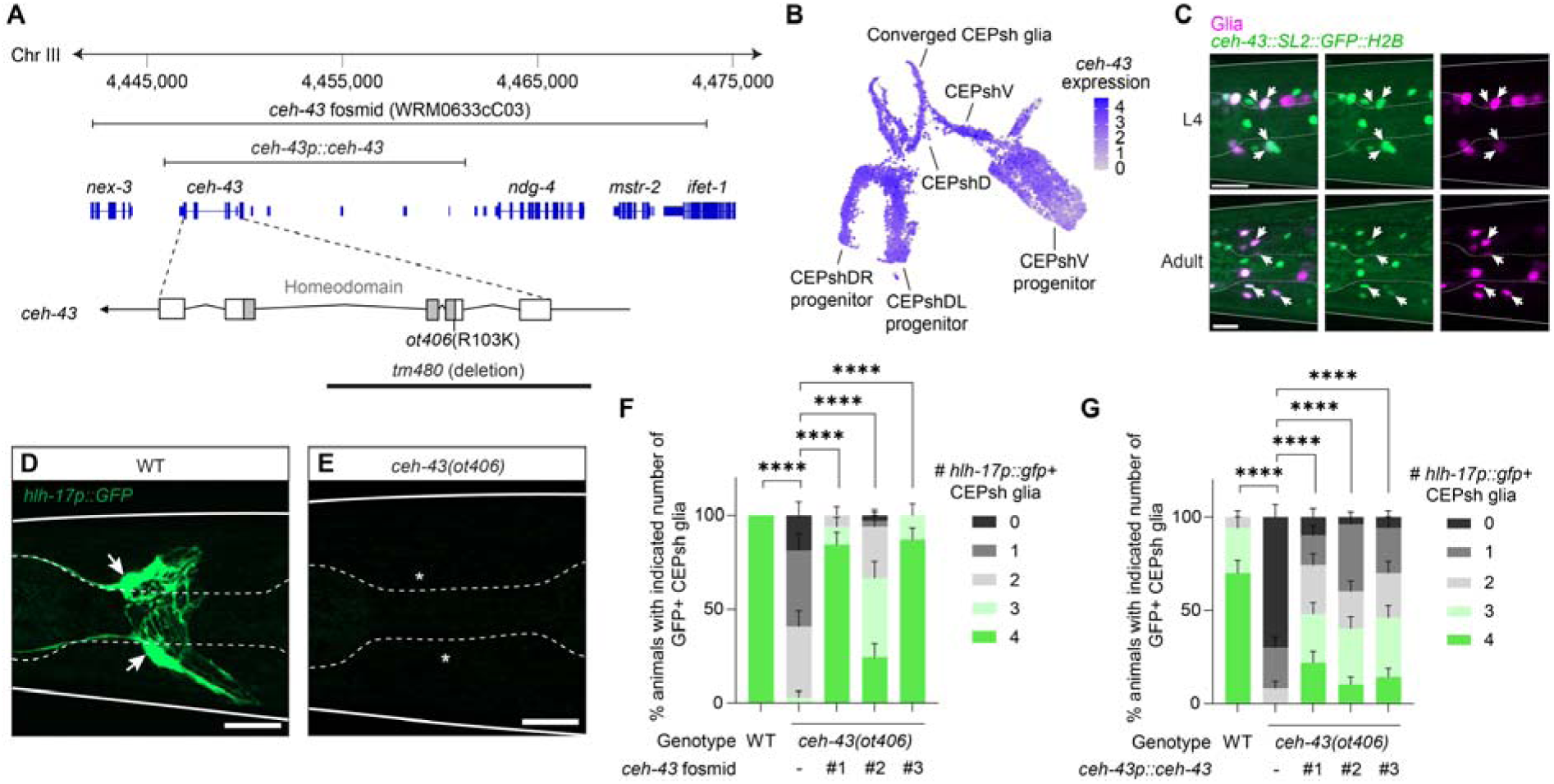
CEH-43 promotes convergent CEPsh glia differentiation. **A**, Schematic of the *ceh-43* locus and DNA constructs used in this study. **B**, Embryonic expression of *ceh-43* in CEPshD and CEPshV lineages. **C**, Post-embryonic *ceh-43* expression (green) in L4 larvae and young adults. Glia (magenta) are labeled with *mir-228p::2xNLS::iBlueberry*. Maximum-projection images of whole animals are shown. Arrows, CEPsh glia nuclei. **D,E**, Representative confocal images of *hlh-17p::gfp* expression in CEPsh glia of L4 larvae in wild-type (WT) (**D**) or *ceh-43(ot406)* mutant (**E**) animals. Arrows, CEPsh glia. Asterisks indicate loss of *hlh-17p::gfp* expression in CEPsh glia. **F**, A *ceh-43* fosmid transgene restores *hlh-17p::gfp* expression to CEPsh glia of *ceh-43*(*ot406*) mutants. n = 31, 32, 32, 33, 31 animals per genotype from three independent scoring sessions. Three independent fosmid rescue lines (#1, #2, #3) were analyzed. \*\*\*\**p* < 0.0001 by Fisher’s exact test. **G**, Same as (**F**) except that rescue was achieved using a PCR fragment containing only *ceh-43* coding and regulatory regions (see panel (**A**)). n = 50 animals per genotype from three independent scoring sessions. Scale bars, 10 μm. Error bars, mean ± s.e.m.

To confirm that CEPsh glia defects arise from *ceh-43* disruption, we introduced wild-type *ceh-43* into *ceh-43(ot406)* mutants. A rescuing fosmid (WRM0633Cc03, **Fig. 3A**) restores *hlh-17p::gfp* expression to mature CEPsh glia in three independent rescue lines (**Fig. 3F**). *hlh-17p::gfp* expression is also restored by a genomic PCR amplicon spanning 10 kb upstream of the start codon and 1.2 kb downstream of the stop codon of the endogenous *ceh-43* locus (*ceh-43p::ceh-43*; **Fig. 3A,G**). These data establish *ceh-43* as the gene required for proper CEPsh glia gene expression, morphology, and differentiation.

### CEH-43 is required for nerve-ring assembly

During embryonic development, the *C. elegans* nerve ring is assembled in a hierarchical sequence in which CEPsh glia first extend processes to demarcate the presumptive neuropil and guide sublateral (SubL) neuron axon extension into it(Rapti et al. 2017). CEPsh glia and SubL neurons then act together to direct follower axons, such as AIY neuron axons(Rapti et al. 2017). CEPsh glia ablation shortly after they are born results in aberrant SubL axon extension and failure of follower neuron nerve-ring entry(Rapti et al. 2017).

We found that c*eh-43(ot406)* mutants phenocopy CEPsh glia ablation. In wild-type animals, SubL neuron axons labeled with *ceh-24p::gfp* enter the nerve ring and extend posteriorly (**Fig. 4A**). In c*eh-43(ot406)* mutants, however, SubL axons exhibit aberrant anterior processes (**Fig. 4B,C, Supplemental Fig. S8A,B**) and other defects (**Supplemental Fig. S8C-G**). Likewise, AIY axons fail to enter the nerve ring in c*eh-43(ot406)* mutants (**Fig. 4D,E, Supplemental Fig. S8H-L**). These defects are rescued by a 14-kb *ceh-43* genomic fragment, confirming that the *ceh-43* lesion is responsible for the nerve-ring defects observed (**Fig. 4F**). Notably, published scRNA-seq data(Packer et al. 2019) and our scRNA-seq data show no detectable *ceh-43* expression in SubL or AIY neurons. Together, these observations indicate that CEH-43 is required in CEPsh glia to orchestrate assembly of the nerve ring.

**Figure 4.**
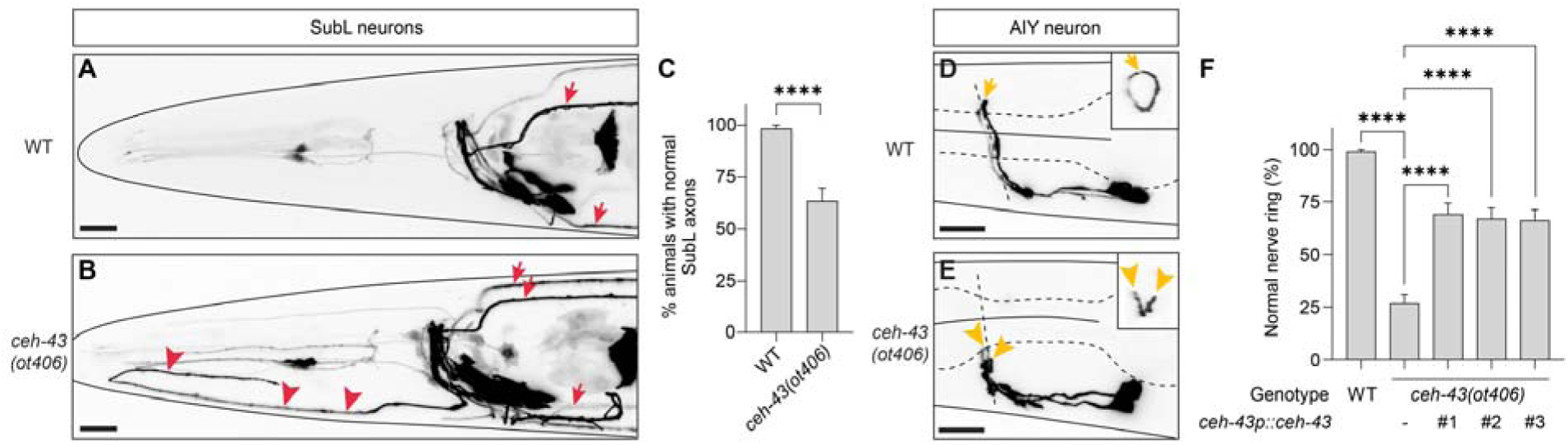
CEH-43 is required for nerve ring assembly. **A,B**, Representative maximum-projection confocal images of sublateral (SubL) neurons labeled with *ceh-24p::gfp* in wild-type (**A**) and *ceh-43(ot406)* (**B**) L4 larvae. Arrows, normal posteriorly extending processes. Arrowheads, aberrant anterior processes. **C**, Quantification of SubL axon guidance defects. n = 60 animals per genotype from three independent imaging sessions. \*\*\*\**p* < 0.0001 by Fisher’s exact test. **D,E**, Representative maximum-projection confocal images of AIY interneurons labeled with *ttx-3p::gfp* in wild-type (**D**) and *ceh-43(ot406)* (**E**) L4 larvae. Arrows, the dorsal apex of left and right AIY axon trajectories. Arrowheads, abnormal AIY axons. **F**, Quantification of AIY axon defects. n = 100, 112, 81, 88, and 86 animals per genotype from three independent imaging sessions. Three independent transgenic lines were analyzed (#1, #2, #3). \*\*\*\**p* < 0.0001 by Fisher’s exact test. Scale bar, 10 μm. Error bars, mean ± s.e.m.

### CEH-43 functions cell-autonomously in early CEPsh glia

To test if CEH-43 functions cell-autonomously to control CEPsh glia differentiation, we generated *ceh-43(ot406)*; *hlh-17p::gfp* mutants carrying an unstable extrachromosomal array consisting of a *ceh-43* fosmid and a ubiquitous *nhr-2p::H1-mCherry* reporter that allows tracking of cells inheriting the fosmid (**Fig. 5A**). Among CEPsh glia carrying the array (mCherry+), 86% (96/111) express *hlh-17p::gfp* (**Fig. 5B,D**). Importantly, all CEPsh glia expressing *hlh-17p::gfp* carry the rescuing fosmid (mCherry+; 96/96; **Fig. 5B,E**), whereas cells that do not express *hlh-17p::gfp* lack the array (mCherry-; 0/6; **Fig. 5C,F**). Similar results are observed with the *ceh-43(tm480)* deletion allele (**Fig. 3A, 5G-I**) or with a pan-glia *mir-228* promoter::*ceh-43* cDNA extrachromosomal transgene (**Supplemental Fig. S7F-N**). Because *mir-228* is expressed in CEPsh glia but not their progenitors (**Supplemental Fig. S7O**), these data indicate that CEH-43 acts in CEPsh glia and not in closely related neurons, epithelial cells, or progenitors.

**Figure 5.**
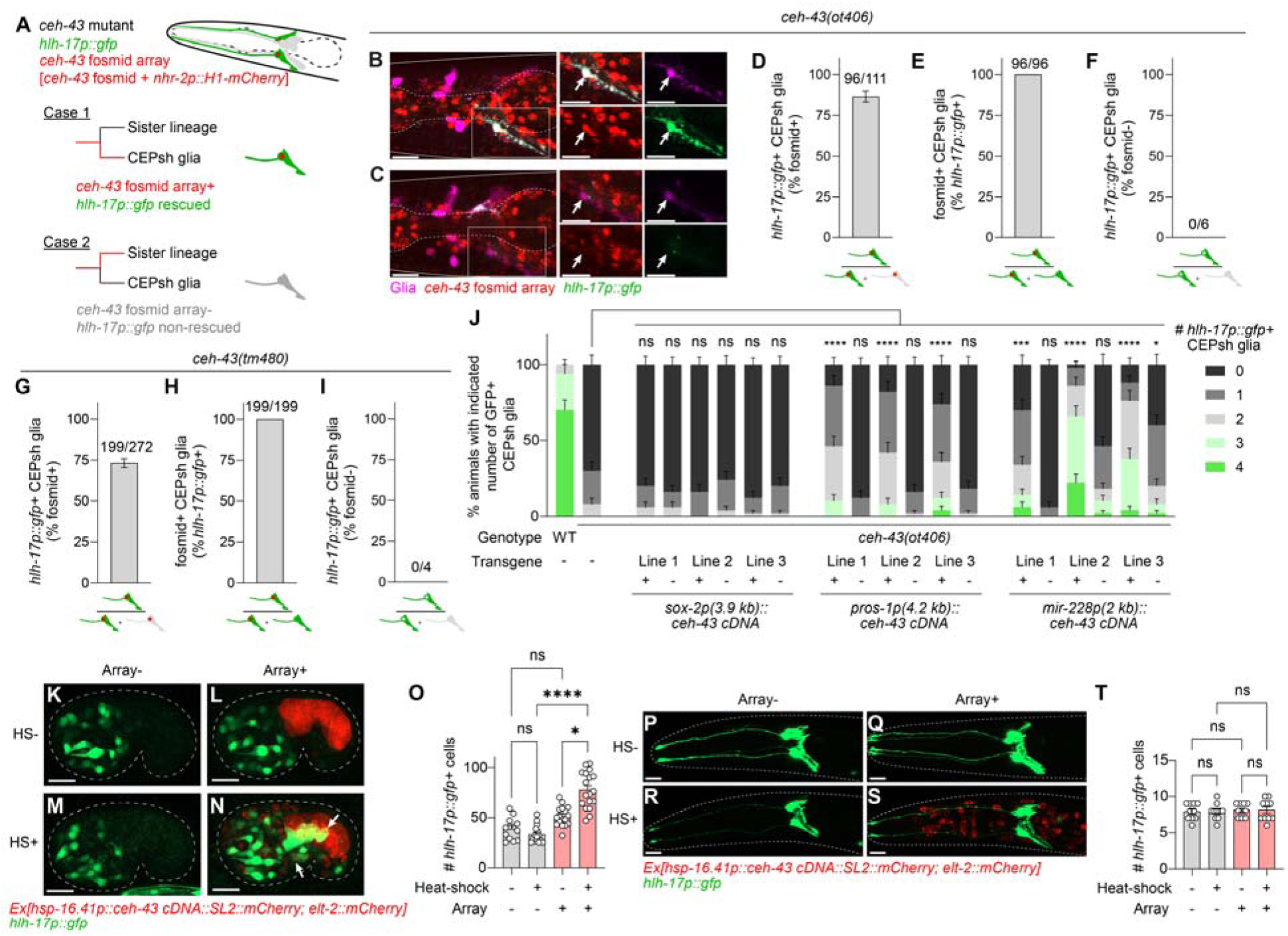
CEH-43 functions cell-autonomously in early CEPsh glia and is sufficient to induce embryonic CEPsh glia gene expression. **A**, General strategy for mosaic analysis. An unstable extrachromosomal array containing wild-type *ceh-43* genomic sequences (fosmid) and a pan-cellular nuclear reporter transgene (*nhr-2p::H1-mCherry*) was introduced into *ceh-43(ot406)* or *ceh-43(tm480)* mutants and assessed for co-expression with the CEH-43 target reporter *hlh-17p::gfp*. If CEH-43 acts cell-autonomously, mCherry-positive CEPsh glia should express GFP (Case 1), whereas mCherry-negative CEPsh glia should not (Case 2). **B**, Presence of the *ceh-43* fosmid, visualized by *nhr-2p::H1-mCherry* (red), overlaps with *hlh-17::gfp* expression (green). Glia are marked with *mir-228p::2xNLS::iBlueberry* (magenta). Insets show magnified views of boxed regions. Arrows indicate CEPsh glia. **C**, Same as (**B**) except that the indicated CEPsh glia (arrow) lacks *nhr-2p::H1-mCherry* (red) and does not express *hlh-17::gfp* expression (green). **D-F**, Quantification of mosaic analysis in *ceh-43(ot406)* mutants. (D) Overall efficiency of *ceh-43* fosmid-mediated restoration of *hlh-17::gfp* expression. (E) All CEPsh glia expressing *hlh-17::gfp* carry the *ceh-43* fosmid. (F) No CEPsh glia lacking the *ceh-43* fosmid express *hlh-17p::gfp*. Numbers above bars indicate total CEPsh glia analyzed across three imaging sessions (30 L4 larvae total). **G-I**, Same as (**D-F**), but for *ceh-43(tm480)* mutants. A total of 70 L4 larvae were examined. **J**, Quantification of rescue efficiency for the indicated transgenes in *ceh-43(ot406)* mutants. n = 50 animals per genotype from >3 independent scoring sessions. \**p* < 0.05, \*\*\**p* < 0.001, \*\*\*\**p* < 0.0001 by Fisher’s exact test. **K-O**, Ectopic CEH-43 expression induces ectopic *hlh-17p::gfp* expression in comma-stage embryos. (**K-N**) Maximum projections of confocal z-stacks. Without heat shock (**K,M**), no ectopic expression is observed in embryos lacking the *hsp-16.41p::ceh-43 cDNA::SL2 mCherry* array (Array-; coinjected with *elt-2::mCherry*, red). HS-Array+ embryos (**L**) display mild ectopic *hlh-17p::gfp* expression, likely due to promoter leakiness, while HS+ Array+ embryos (**N**) exhibit stronger ectopic expression. (O) Quantification of the heat-shock experiment. n = 15, 13, 18, and 17 embryos from 2–3 imaging sessions. \**p* < 0.05; \*\*\*\**p* < 0.0001; ns, not significant by Kruskal-Wallis test with Dunn’s multiple comparisons correction. (**P-T**) Same as (**K-O**), except performed in L4 larvae. (**T**) n = 10 animals from two imaging sessions. Scale bars, 10 μm. Error bars, mean ± s.e.m.

Further supporting this conclusion, *pros-1* is expressed in both CEPsh glia and their progenitors (**Supplemental Fig. S7D**), and a *pros-1*p::*ceh-43* cDNA transgene, containing 4.2 kb of *pros-1* upstream regulatory sequences, can partially but significantly rescue *ceh-43(ot406)*; *hlh-17p::gfp* mutants (**Fig. 5J**). Rescue is also observed using a transgene with a 2 kb *mir-228* promoter fragment (**Fig. 5J**). By contrast, *sox-2* is expressed primarily in CEPsh glia progenitors (**Supplemental Fig. S7E**), and a *sox-2*p::*ceh-43* cDNA transgene containing 3.9 kb of upstream regulatory sequences fails to restore *hlh-17p::gfp* expression to *ceh-43(ot406)* mutants (**Fig. 5J**).

Thus, our data support a model in which CEH-43 acts cell autonomously within early CEPsh glia to promote their differentiation, rather than acting in progenitors or neighboring non-glial cells.

### CEH-43 expression is sufficient to induce embryonic CEPsh glia gene expression

CEH-43 could function permissively, allowing other cell-specific regulators to activate CEPsh glia gene expression, or it may directly promote CEPsh glia gene expression. To distinguish between these possibilities, we ectopically expressed *ceh-43* cDNA using a heat-shock inducible *hsp-16.41* promoter in embryos carrying the *hlh-17p::gfp* reporter transgene (**Fig. 5K-N**). Even without heat shock, *hsp-16.41p::ceh-43* transgenic embryos show a modest increase in the number of *hlh-17p::gfp*-positive cells, likely due to leaky transgene expression (**Fig. 5L,O**). Upon heat shock, the number of ectopic *hlh-17p::gfp*-positive cells further increases (**Fig. 5N,O**), indicating that CEH-43 is sufficient to activate embryonic CEPsh glia gene expression.

Heat-shock induction of *ceh-43* expression in L4 larvae fails to induce ectopic *hlh-17p::gfp* expression (**Fig. 5P-T**), suggesting that CEH-43 may only be able to activate CEPsh glia gene expression in embryos. To test whether CEH-43 is nonetheless required at later stages to maintain CEPsh glia gene expression, we generated animals expressing an endogenously encoded CEH-43::3xGAS::GFP-AID (<u>a</u>uxin-inducible <u>d</u>egron(Nishimura et al. 2009; Zhang et al. 2015)) protein and a ubiquitously expressed TIR1 E3 ligase (*eft-3p::TIR1-mRuby*), enabling auxin-induced protein degradation (**Supplemental Fig. S9A**). Treatment of early L1 stage larvae with 4 mM auxin completely degrades CEH-43::3xGAS::GFP::AID within two hours (**Supplemental Fig. S9B**) but does not affect *hlh-17p::gfp* expression in CEPsh glia (**Supplemental Fig. S9C-G**). Similarly, auxin-induced degradation of CEH-43 has little effect on larval expression of the CEPsh glia gene *glt-1* (**Supplemental Fig. S9H-L**).

These findings indicate that CEH-43 is required during embryogenesis to establish CEPsh glia gene expression but is largely dispensable for its maintenance. Alternatively, CEH-43 may maintain the expression of some CEPsh glia genes, but not others.

### CEH-43 binds genes expressed during both phases of CEPsh glia development

To determine how CEH-43 controls CEPsh glia development, we identified its *in vivo* targets using NanoDam(Tang et al. 2022) (**Fig. 6A**). We generated animals in which *gfp* coding sequences are endogenously fused to the *ceh-43* gene. These animals also carry an *hsp-16.2*p::Dam-GFP Nanobody transgene, containing a fusion of *E. coli* Dam methyltransferase to GFP nanobody sequences (modified from (Gómez-Saldivar et al. 2020)). Binding of the Dam-GFP Nanobody to DNA-bound CEH-43::GFP promotes methylation of nearby adenine residues in 5’-GATC-3’ motifs, and mapping these methylated sites enables identification of CEH-43 binding loci. Because *C. elegans* genomic DNA is normally unmethylated(Greer et al. 2015), the approach provides a high signal-to-noise ratio. Importantly, the permanent methylation signature allows simultaneous identification of embryonic and postembryonic CEH-43 binding sites when L4 larvae are interrogated. Two L4 biological replicates we collected show strong concordance (**Supplemental Fig. S10**).

**Figure 6.**
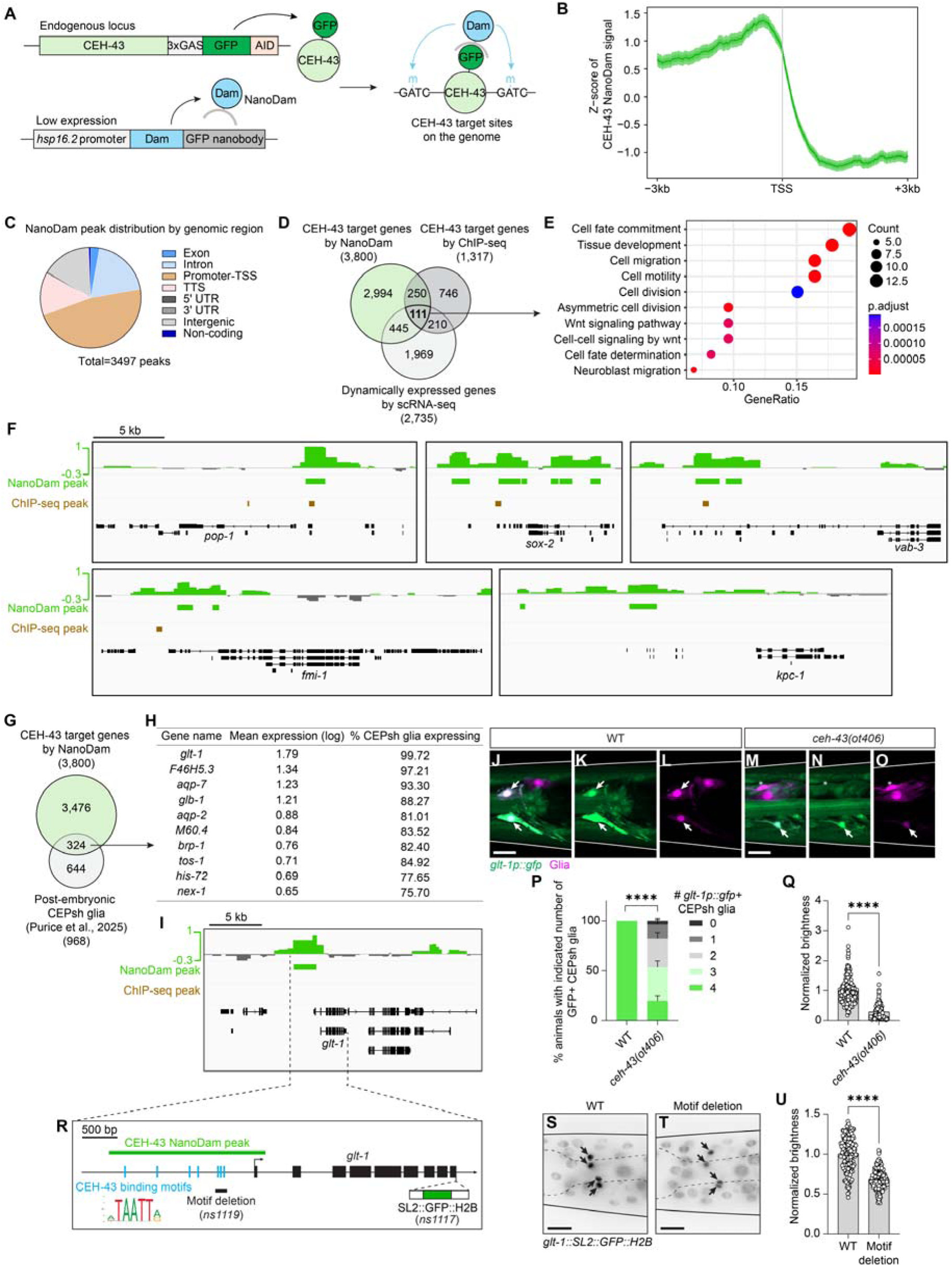
CEH-43 binds genes expressed during both phases of CEPsh glia development. **A**, Schematic of the CEH-43 NanoDam strategy. A leaky *hsp-16.2* promoter drives low-level expression of a GFP nanobody-Dam methylase fusion protein (NanoDam) in all cells without heat shock. Endogenous CEH-43 protein is C-terminally tagged with GFP. The GFP nanobody binds GFP-tagged CEH-43, recruiting Dam methylase to CEH-43-occupied DNA binding sites, leading to methylation of nearby GATC sequences. **B**, Aggregation plot of CEH-43 NanoDam signal (Z-score) centered on transcription start sites (TSS) across all detected peaks, showing enrichment upstream of TSS. **C**, Genomic distribution of 3,497 NanoDam peaks, revealing enrichment in non-coding regions. Peaks were called at a false discovery rate (FDR) <10^-25^. TTS, transcription termination site. **D**, Venn diagram showing overlap between CEH-43 target genes identified by NanoDam, ChIP-seq (Kudron et al., 2024), and dynamically expressed genes in the CEPsh glia lineage from the scRNA-seq dataset (Fig. 1). **E**, Gene Ontology (GO) enrichment analysis of 111 high-confidence embryonic CEH-43 target genes. **F**, Representative CEH-43-targeted genes. NanoDam binding intensities are shown as log_2_-fold enrichment of CEH-43 NanoDam over NanoDam-alone controls. Identified binding peaks are marked in green. Corresponding CEH-43 ChIP-seq peaks from Kudron et al. (2024) are shown in brown. Gene names and exon-intron structures are shown below. **G**, Overlap between CEH-43 NanoDam targets and genes expressed in post-embryonic CEPsh glia (Purice et al., 2025). **H**, Top 10 CEH-43 NanoDam targets ranked by expression level in post-embryonic CEPsh glia (Purice et al., 2025). **I**, NanoDam binding profile upstream of *glt-1* coding region. **J-O**, CEH-43 is required for *glt-1* expression. Maximum-projection confocal images of representative wild-type (**J-L**) and *ceh-43(ot406)* (**M-O**) L4 larvae. (**J,M**) Merged channels; (**K,N**) *glt-1p::gfp*; (**L,O**) *mir-228p::2xNLS::iBlueberry* pan-glial marker. Arrows, *glt-1p::gfp*-positive CEPsh glia. Asterisks, CEPsh glia lacking *glt-1p::gfp* expression. (P) Quantification showing significantly fewer *glt-1p::gfp*-positive CEPsh glia in *ceh-43(ot406)* mutants compared with wild type. \*\*\*\**p* < 0.0001 by Fisher’s exact test. (Q) Quantification of *glt-1p::gfp* fluorescent intensity in CEPsh glia, showing significant reduction in *ceh-43(ot406)* mutants compared with wild type. \*\*\*\**p* < 0.0001 by Welch’s t-test. (R) Schematic showing predicted CEH-43 binding motifs (blue) within the CEH-43 NanoDam peak (green) upstream of the *glt-1* transcription start site. (**S,T**) Representative maximum-projection images of WT animals and animals with CEH-43 motif deleted. CEPsh glia are indicated by arrows. (**U**) Quantification of *glt-1::SL2::GFP::H2B* brightness in CEPsh glia. \*\*\*\**p* < 0.0001 by Welch’s t-test. Scale bar, 10 μm. Error bars, mean ± s.e.m.

Consistent with a role in transcriptional control, CEH-43 binding peaks (called at a false discovery rate (FDR) <10^-25^, see **Methods**)(Tang et al. 2022) are enriched upstream of transcription start sites (TSS; **Fig. 6B**). 97% of binding peaks are located in non-exonic regions, even though these regions comprise only 75.8% of the genome and are less likely to contain GATC motifs than exons (predicted ratio of 1:1.39(Amit et al. 2012); **Fig. 6C**). Moreover, 361 target genes identified using NanoDam overlap with CEH-43 chromatin immunoprecipitation sequencing (ChIP-seq) targets identified in a previously published study of mixed-stage embryos(Kudron et al. 2024) (**Fig. 6D**).

Cross referencing NanoDam predicted CEH-43 binding sites with our embryonic CEPsh glia scRNA-seq dataset identifies 111 high-confidence embryonic CEH-43 targets (**Fig. 6D, Supplemental Table S6**). Gene ontology analysis reveals enrichment for genes involved in asymmetric cell division, fate commitment, migration, and Wnt signaling (**Fig. 6E**). These include *pop-1*, a Wnt signaling effector regulating asymmetric cell division(Mizumoto and Sawa 2007); *sox-2*, a regulator of neuronal fate(Alqadah et al. 2015); and *vab-3*, involved in CEPsh glia specification(Yoshimura et al. 2008) (**Fig. 6F**). Other target genes include *fmi-1*, Flamingo/CELSR/ADGRC adhesion GPCR; and *kpc-1*, a Furin-like protease, both required in CEPsh glia to promote nerve-ring axon extension(Rapti et al. 2017) (**Fig. 6F**). These results indicate that CEH-43 controls expression of genes during phase I of CEPsh glia development (**Fig. 2J**).

To determine whether CEH-43 also binds genes expressed during phase II, we compared NanoDam-identified targets with genes expressed post-embryonically in CEPsh glia(Purice et al. 2025). 324 putative postembryonic targets were identified (**Fig. 6G, Supplemental Table S7**). Among these, *glt-1*, encoding a glutamate transporter homologous to mammalian GLT1/EAAT2, is highly expressed (**Fig. 6H,I**). Importantly, expression of a *glt-1p::gfp* reporter in CEPsh glia is significantly reduced in *ceh-43(ot406)* mutants (**Fig. 6J-Q**). Deleting three predicted CEH-43 binding motifs(Catron et al. 1993; Doitsidou et al. 2013) within the CEH-43 NanoDam peak significantly reduces expression of the endogenous *glt-1::SL2::GFP::H2B* reporter (**Fig. 6R-U**). Thus, CEH-43 also controls gene expression during phase II of CEPsh glia development (**Fig. 2J**).

*glt-1* was not identified as a CEH-43 target in previous embryonic ChIP-seq dataset(Kudron et al. 2024), nor is it expressed in our converged early embryonic CEPsh glia dataset. Furthermore, *glt-1p::gfp* is already expressed in early L1 larvae, and auxin-induced depletion of CEH-43 starting at the L1 stage has little effect on *glt-1p::gfp* reporter expression (**Supplemental Fig. S9H-L**). We suggest, therefore, that CEH-43 acts during late embryonic or very early postembryonic stages to initiate *glt-1* expression. More broadly, these findings suggest that the second phase of CEPsh glia maturation (**Fig. 2J**) is initiated in later embryos or in very early L1 animals.

Finally, we note that *eat-6*, homologous to the mammalian *ATP1A2* gene encoding the astrocytic Na^+^/K^+^-ATPase α2 subunit, is co-expressed with *glt-1* in adult CEPsh glia(Purice et al. 2025) and is bound by CEH-43 (**Supplemental Fig. S10C**). In mammals, α2 is essential for maintaining astrocytic Na^+^ homeostasis, which in turn supports GLT1- and GLAST-mediated glutamate uptake(Melone et al. 2019; Cholet et al. 2002; Rose et al. 2009). Loss of α2 impairs glutamate clearance and is linked to cortical dendritic hyperexcitability and cranial pain in a mouse model of familial hemiplegic migraine(Romanos et al. 2020). The co-expression and CEH-43-dependent regulation of *eat-6* and *glt-1* suggests conserved mechanisms of astrocytic glutamate regulation from *C. elegans* to mammals.

### Gene expression similarities between *C. elegans* and mammalian astrocytes

Developmental, morphological, and functional parallels between *C. elegans* CEPsh glia and mammalian radial glia and astrocytes(Nagai et al. 2021) raise the possibility that these cell types also share conserved transcriptional signatures. To investigate this, we re-analyzed published scRNA-seq datasets of cultured human glial progenitor cells, human induced pluripotent stem cells (iPSC), and mouse embryonic stem cells (mESCs) undergoing differentiation(Wang et al. 2025; Frazel et al. 2023) (**Fig. 7A**; Wang et al. 2025, Frazel et al. 2023 (Human), and Frazel et al. 2023 (Mouse), respectively). Using criteria from the original studies, we identified radial glia, astrocyte, and neuronal clusters in each dataset (**Supplemental Fig. S11A-F**). For each dataset, we identified the top 50 radial-glia-, astrocyte-, or neuron-enriched genes with *C. elegans* orthologs (**Supplemental Table S8-S10**) and calculated their normalized average expression across CEPsh glia lineages (**Fig. 7A**, bottom). In the Wang et al. 2025 dataset, orthologs of radial-glia-enriched genes are comparably expressed in early CEPshD, early CEPshV, and converged CEPsh glia, and are expressed at higher levels than in progenitors and URA neurons, consistent with early CEPsh glia taking on a radial-glia-like state (**Fig. 7B**). Astrocyte-enriched orthologs are most highly expressed in converged CEPsh glia, with moderate expression in nascent CEPsh glia and minimal expression in progenitors, also consistent with converged CEPsh glia transitioning along a radial-glia-to-astrocyte developmental trajectory. By contrast, neuron-enriched orthologs show the opposite pattern: strongly expressed in URA neurons but significantly less in converged CEPsh glia (**Fig. 7B**). The human iPSC (Frazel et al. 2023 (Human)) and mouse ESC (Frazel et al. 2023 (Mouse)) datasets show similar overall trends (**Fig. 7B-D**).

**Figure 7.**
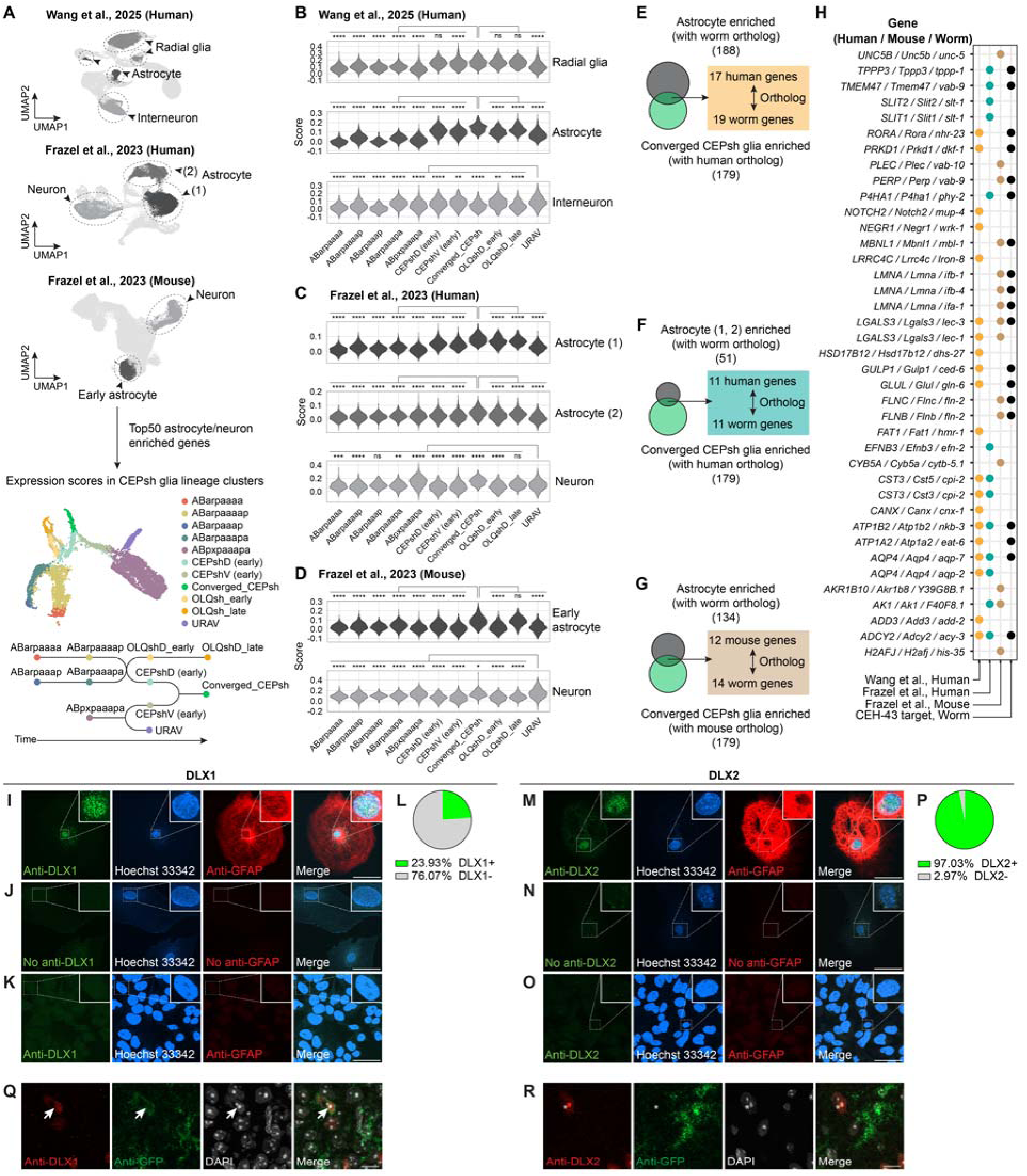
Conserved transcriptional and molecular features of *C. elegans* and mammalian astrocytes. **A**, Top, UMAP projections of three published single-cell RNA-seq datasets (Wang et al., 2025; Frazel et al., 2023, human; Frazel et al., 2023, mouse) highlighting radial glia, astrocyte, and neuron populations. Bottom, UMAP projection of cell types across the *C. elegans* CEPsh glia lineage, with a schematic of the lineage relationships. **B-D**, Module scores of radial glia-, astrocyte-, and neuron-enriched genes from the Wang et al. (2025) (**B**), the Frazel et al. (2023) human (**C**), and the Frazel et al. (2023) mouse (**D**) datasets, plotted across *C. elegans* CEPsh lineage clusters. Statistical significance was determined by Wilcoxon test with Bonferroni correction. \**p* < 0.05; \*\**p* < 0.01; \*\*\*\**p* < 0.0001, ns, not significant. **E-G**, Orthologous gene pairs enriched in both CEPsh glia and mammalian astrocytes (average log_2_FC >0.5, adjusted *p*-value <0.05): (**E**) Wang et al. (2025); (**F**) Frazel et al. (2023, human); (**G**) Frazel et al. (2023, mouse). **H**, Genes enriched in both CEPsh glia and mammalian astrocytes. Dot plot showing the presence of each gene in human and mouse astrocyte datasets and whether the corresponding *C. elegans* ortholog is a CEH-43 NanoDam target. **I**, Immunofluorescence staining of primary astrocytes showing DLX1, GFAP, and Hoechst 33342 nuclear counterstain. Representative Z-stack images are shown. Scale bar, 50µm. **J**, Same as (**I**) but without primary antibodies (negative control). **K**, Same as (**I**) but staining undifferentiated P19 cells (negative control). **L**, Quantification of the proportion of DLX1+ astrocytes. **M**, Immunofluorescence staining of primary astrocytes showing DLX2, GFAP, and Hoechst 33342. Representative Z-stack images are shown. Scale bar, 50µm. **N**, Same as (**M**) but without primary antibodies (negative control). **O**, Same as (**M**) but staining undifferentiated P19 cells (negative control). **P**, Quantification of the proportion of DLX2+ astrocytes. **Q**, Immunofluorescence staining of adult JD130 (*Aldh1l1-EGFP/Rpl10a*) mouse brain showing a DLX1+ astrocyte (arrow) in the retrosplenial cortex. Representative Z-stack images are shown. Scale bar, 10µm. **R**, Same as (**Q**) but showing a DLX2+ non-astrocyte (asterisk).

We also compared the entire list of astrocyte-enriched genes from each of the three mammalian datasets with genes enriched in converged CEPsh glia (**Fig. 7E-G, Supplemental Table S11**). The combined set of overlapping orthologs (**Fig. 7H**) includes sodium/potassium-transporting ATPase subunits (*eat-6/ATP1A2*, *nkb-3/ATP1B2*), cell adhesion molecules (*lec-1/lec-3/LGALS3*, *wrk-1/NEGR1*), intercellular signaling components (*slt-1/SLIT1/2*), GFAP-related intermediate filament genes (*ifa-1*, *ifa-4*, *ifb-1*), and aquaporins (*aqp-2,7/AQP4*), among others. Notably, many of these genes also contain CEH-43 NanoDam binding peaks, suggesting that CEH-43 regulates a conserved core astrocytic gene expression module (**Fig. 7H**).

Together, these analyses indicate that despite their evolutionary distance, mammalian and *C. elegans* radial glia and astrocytes share recognizable transcriptional features.

### The CEH-43 homologs DLX1 and DLX2 are expressed in mammalian astrocytes

Of 327 transcription factors expressed in CEPsh glia or their progenitors, 201 have at least one identifiable mouse or human orthologs. Among 671 putative human homologs, 323 and 318 are expressed in the Wang et al.(Wang et al. 2025) and Frazel et al.(Frazel et al. 2023) human astrocyte datasets, respectively. Similarly, 317 of 551 putative mouse orthologs are expressed in the Frazel et al. mouse dataset(Frazel et al. 2023) (**Supplemental Fig. S11G, Supplemental Table S12**). Shared factors included *NFIA*, a regulator of astrocyte generation(Deneen et al. 2006; Kang et al. 2012; Tchieu et al. 2019); *TCF7L2*, involved in astrocyte maturation(Szewczyk et al. 2024); and *PROX1/prospero*, a regulator of glia development in *Drosophila* and mice(Freeman and Doe 2001; Stergiopoulos et al. 2015) (**Supplemental Fig. S11H**). Although *C. elegans* lacks a direct *SOX9* ortholog, related *Sox* family members (*sox-2* and *sem-2*) are expressed in CEPsh progenitors, and SOX-2 is the most similar *C. elegans* protein to mammalian *SOX9*.

Importantly, we also found that both DLX1 and DLX2, orthologs of *C. elegans* CEH-43, are expressed in subsets of astrocytes (**Supplemental Fig. S11H**). This was unexpected given well-established roles for these transcription factors in dopaminergic and GABAergic neuron differentiation(Anderson et al. 1997a; Andrews et al. 2003; Brill et al. 2008; Le et al. 2017). To directly test whether mammalian astrocytes express DLX proteins, we immuno-stained fixed primary astrocytes isolated from embryonic mouse brain. Remarkably, both DLX1 and DLX2 are detected, and their localization is predominantly nuclear, consistent with their functions as transcription factors. (**Fig. 7I-P**). We also examined DLX1 and DLX2 expression *in vivo*. In the retrosplenial cortex of adult JD130 (*Aldh1l1-EGFP/Rpl10a*) mice, DLX1 but not DLX2 is detected in astrocyte nuclei (**Fig. 7Q,R**). These findings suggest that, like CEH-43 in *C. elegans*, DLX1 and DLX2 may regulate astrocyte gene expression during early mammalian astrocyte development. Consistent with the mammalian gene expression datasets, neither DLX1 nor DLX2 is expressed in all astrocytes (**Supplemental Fig. S11H**).

## Discussion

Our studies highlights the power of lineage-restricted single-cell transcriptomics for resolving cell-type-specific developmental trajectories and allow us to assemble a high-resolution transcriptional roadmap for early development and cell-division-independent radial-glia-to-astrocyte transformation of *C. elegans* CEPsh glia. Similar trans-differentiation is common in vertebrate brain development(Voigt 1989; Brunne et al. 2010; Tabata 2015; Culican et al. 1990), and our findings suggest molecular conservation of the process across species. We show that CEPsh glia develop through two distinct phases (**Fig. 2J**). First, nascent dorsal and ventral CEPsh glia rapidly converge onto a shared transcriptional state enriched for cell adhesion and axon guidance genes. CEPsh glia bipolar morphology and pioneer roles in nerve-ring assembly at this stage(Rapti et al. 2017) strongly suggest similarity to vertebrate radial glia, a proposition supported by gene expression similarities. In the second phase, which likely begins in later embryos, after nerve-ring assembly, CEPsh glia develop elaborate processes, ensheathe synapses, and acquire a distinct expression profile enriched for genes involved in synaptic and neuronal activity(Katz et al. 2019; Purice et al. 2025), resembling vertebrate astrocytes.

The mechanisms underlying direct radial-glia-to-astrocyte transformation in vertebrates remain largely unknown. Our findings, however, suggest that key aspects of this process may be conserved. We identify the *distal-less* transcription factor CEH-43 as a key CEPsh glia factor governing gene expression in both the radial glia and astrocyte forms of these cells. Roles for the murine CEH-43 homologs DLX1/2 in radial glia were previously described in the context of dopaminergic(Andrews et al. 2003; Brill et al. 2008) and GABAergic(Anderson et al. 1997a; Le et al. 2017) neuron specification. We now show that these genes, like CEH-43, are also expressed in astrocytes *in vivo*, in line with published scRNA-seq and histological datasets(Allen et al. 2022a; Uhlén et al. 2015). While DLX gene expression appears widespread in fetal astrocytes, we detect many fewer DLX expressing astrocytes in adults. One possible explanation for this may be that in adulthood, most radial glia have already converted to astrocytes, so that fewer such transitions can be detected.

Our studies also reveal that CEH-43 mediates the convergent differentiation of CEPsh glia that emerge from lineally and transcriptionally distinct progenitors. Similar instances of transcriptional convergence have also been described in mammalian settings. Oligodendrocyte precursor cells, for example, are generated from spatially distinct regions of the neural tube but converge onto a shared transcriptional state characterized by *Sox10* expression(Zeisel et al. 2018; Marques et al. 2018). Astrocytes derived from different radial glial domains also converge onto a common transcriptional state(Bandler et al. 2022). How convergent differentiation is regulated in these settings is largely unknown. In *C. elegans*, cholinergic neurons derived from different lineages express a shared suite of genes regulated by a single transcription factor, UNC-3(Kratsios et al. 2012). CEH-43 may have similar roles in CEPsh glia. However, we previously reported that the MLS2/HMX transcription factor preferentially affects ventral CEPsh glia gene expression(Yoshimura et al. 2008), suggesting that lineage-selective transcription factors(Mizeracka et al. 2021) also contribute to CEPsh glia specification. Furthermore, whereas terminal selector transcription factors like UNC-3 initiate and maintain cell-type-specific gene expression(Hobert 2008; Hobert and Kratsios 2019; Stefanakis et al. 2024, 2025), and CEH-43 functions similarly in dopaminergic neurons(Doitsidou et al. 2013), CEH-43 is not required to maintain CEPsh glia gene expression, and its ectopic expression induces target gene expression only during embryogenesis. Thus, in CEPsh glia, CEH-43 appears to act only as an early instructive regulator that drives gene expression when terminal gene expression is first established. Future studies will determine whether this mode of action is also applicable to mammalian DLX factors.

## Materials and Methods

### C. elegans husbandry

*C. elegans* were maintained at 20°C on NGM agar plates supplemented with cholesterol and seeded with OP50 *E.coli* strain as food, following standard protocols(Brenner 1974; Stiernagle 2006).

### Mouse housing and procedures

All experiments using mice were reviewed and approved by the Rockefeller University IACUC and adhere to the institutional and NIH rules and regulations. We utilized two mouse strains purchased from the Jackson Laboratory: C57Bl/6J mice (JAX strain # 000664) and astrocyte specific TRAP mouse line in which L10a-GFP is expressed under the astrocyte specific *Aldh1l1* promoter(Doyle et al. 2008), *b6;fvb-tg(aldh1l1-egfp/rpl10a)jd130htz/j* (JAX strain # 030247). Mice were group housed in conventional rooms in a 12-hour light/dark cycle and access to food and water ad libitum. Mice were perfused by transcardial perfusion at 20-24 weeks. Prior to transcardial perfusion and tissue harvesting, mice were anesthetized with a lethal intraperitoneal injection of ketamine (300 mg/kg in saline).

### Preparation of synchronized embryos for single-cell dissociation

*C. elegans* embryos were harvested using a standard hypochlorite bleaching protocol(Stiernagle 2006) and incubated in M9 buffer(Stiernagle 2006) without food until L1 larvae hatched. L1s were then plated onto standard 150 mm NGM plates seeded with *E. coli* OP50, with no more than 15,000 L1s per plate, and raised at 20°C. Plates were monitored regularly, and concentrated OP50 was added as needed when food became scarce. After 48–48.5 hours (timing may vary by strain), young adult animals were checked under a dissecting microscope to assess embryo content. Three fields of view from each of three plates were examined. When 20–30% of animals had a single embryo in one arm of the gonad, the population was collected via hypochlorite bleaching. The time of hypochlorite treatment was designated as t = 0. Embryos that survived the bleach treatment were washed three times with M9 and once with egg buffer(Zhang and Kuhn 2018), then brought to 2 mL total volume in a 15 mL Falcon tube. To remove debris, embryos were purified via one round of sucrose density gradient centrifugation: 4 mL of 50% sucrose solution was added, vortexed to mix, and centrifuged at 450 g for 3 minutes. A layer of pure embryos was collected from the top, while debris and animal carcasses were pelleted at the bottom. Embryos were transferred with a glass pipette to a fresh 15 mL tube and washed three times with egg buffer (fill to 15 mL, centrifuge at 450 g for 3 minutes). After the final wash, embryos were resuspended in 10 mL egg buffer and grown on a rotor at 20°C.

### Single-cell dissociation, FACS, and single-cell RNA sequencing

At the desired developmental stage, embryos were spun down, and the volume was reduced to 0.5 mL. Chitinase (15.6 μL of 20 mg/mL stock; Sigma C8241-25UN) was added to reach a final concentration of 1 U/mL. Embryos were then incubated on a shaker/rotor at 20 °C for 25 minutes. Following digestion, embryos were pelleted with a desk-top centrifuge, the chitinase solution was removed, and 100 μL of freshly prepared pronase (15 mg/mL, Sigma P8811-1G) was added. Embryos were dissociated by repeated passage through a 22-gauge, 1½-inch needle attached to a 3 mL syringe for 2–3 minutes. Dissociation was stopped by adding 1 mL ice-cold egg buffer with 1% BSA. From this point onward, all steps were carried out on ice. The time of ice-cold BSA addition (i.e. cells put on ice) was considered the endpoint (t_end). Dorsal samples were collected at t_end = 240, 270 (two replicates), 330 (two replicates), and 420 minutes. Ventral samples were collected at t_end = 240, 330, and 420 minutes.

Cells were centrifuged at 1,500 g for 5 minutes at 4 °C and washed three times with 1 mL ice-cold egg buffer containing 0.15% BSA. During the final wash, a Millex™-SV 5.0 μm filter was pre-wetted with 1 mL egg buffer with 0.15% BSA. After washing, the cell pellet was resuspended in 500 μL ice-cold egg buffer with 0.15% BSA and passed through the filter into a 1.7 mL low-retention tube (Denville Scientific Inc., Cat. C2170). Residual cells were recovered by rinsing the filter with an additional 0.5–0.6 mL egg buffer into the same tube. Cell density was measured with a hemocytometer and adjusted to ≤1 × 10^7^ cells/mL prior to FACS. DAPI (10 ng/mL, diluted from Thermo Scientific™ Cat. 62248) was used to exclude dead cells during the sorting.

Collection tubes for FACS were pre-coated overnight at 4 °C with egg buffer containing 5% BSA. Target cell populations were sorted into these pre-coated tubes. DAPI (10 ng/mL final concentration) was used for dead cell exclusion. Sorted cells were immediately processed for single cell capture and library preparation using the 10x Genomics Chromium Next GEM Single Cell 3’ Reagent Kits v3.1. Each sample was loaded into a separate channel. For dorsal and ventral samples collected at t_end = 420 minutes, 20,000 cells were loaded. For all other samples, 10,000 cells were loaded.

### Single-cell RNA-seq data processing, quality control, and integration

The *C. elegans* reference genome (WBcel235) was appended with fluorescent marker transgene sequences. For each sample, sequencing reads were aligned to the modified reference genome using 10x Genomics Cell Ranger (v7.1.0) with default settings, including the inclusion of intronic reads. The resulting feature-barcode matrices were processed using the CellBender (v0.2.1) pipeline(Fleming et al. 2022) for ambient RNA removal. Filtered matrices were then imported into Seurat (v4)(Hao et al. 2021) for downstream analysis. Cells with fewer than 200 detected genes (nFeature_RNA ≤ 200) or with >5% mitochondrial RNA content were excluded.

Doublets were identified using the scDblFinder pipeline(Germain et al. 2022) and DoubletFinder(McGinnis et al. 2019). For CEPshD lineages defined based on *pros-1* and *ham-2* expression, doublets were identified by scDblFinder. Clusters dominated by doublets were removed. For remaining clusters, doublets identified by scDblFinder were also removed. For the ventral dataset, doublets were identified with both methods, and clusters with >20% doublet rates according to both methods were excluded. Additional intermediate clusters were also removed if their top enriched genes were predominantly mitochondrial, or if they were located between major lineage branches and consistently co-expressed marker genes from both. Finally, lineages not belonging to targeted lineages shown in **Fig. 1B** were excluded from further analysis.

For the ventral dataset, 2000 integration features were used to perform Seurat (v4) integration to remove batch effects. Integration did not affect the temporal relationships among cells but did eliminate batch differences between samples collected at the same developmental time.

### Cell type annotation and embryo time estimation

Cell lineages and types were annotated based on marker genes curated from previously published datasets(Packer et al. 2019; Ma et al. 2021; Murray et al. 2012; Reilly et al. 2020). For the ventral dataset, only cells from the ABpl/rpaa lineage were retained for analysis. For the dorsal dataset, initial cell type labels were assigned via label transfer using Seurat’s FindTransferAnchors() and TransferData() functions, referencing the VisCello dataset(Packer et al. 2019). Two parallel rounds of label transfer were conducted: one using only ABa lineage cells as the reference, and another using the full VisCello dataset. Final annotations were refined based on these results in conjunction with curated lineage data from the literature.

Embryo time analysis was performed according to a previously published method(Packer et al. 2019), with code generously provided by Dr. John I. Murray. Briefly, gene expression profiles of each single cell were compared to a published whole-embryo gene expression time course(Hashimshony et al. 2015). Time-dependent genes from the reference dataset, defined as those with >0.6 autocorrelation and >1.5 standard deviation in expression, were retained for analysis. For each single cell, genes with log_2_(expression) <1 or >7 were excluded. Pearson correlation coefficients were then calculated against all time points in the reference time course. After applying LOESS(Jacoby 2000) to the correlation profile, the time point with the maximal correlation was assigned as the estimated embryo time.

### Differential Gene Expression and GO Enrichment Analysis

Differential gene expression (DEG) analyses were performed using the FindMarkers() and FindAllMarkers() functions with default settings in Seurat (v4). For heatmap visualization in **Fig. 2B**, the top six differentially expressed transcription factors were selected and supplemented with transcription factors broadly expressed across dorsal and ventral CEPsh glia lineages. These were defined as transcription factors enriched in CEPsh glia lineages by ≥0.5 log_2_FC and expressed in >50% of cells, but not highly differentially expressed within CEPsh glia lineage subclusters, i.e., with all subcluster log_2_FC enrichment <1.3. Converged CEPsh enriched genes in **Fig. 2G** were defined as those with log2FC >1.5 and adjusted p-value (p_val_adj) <0.05. GO enrichment analyses were performed using the enrichGO() function in the clusterProfiler package(Wu et al. 2021; Yu et al. 2012).

To identify dynamically expressed genes in CEPsh glia lineages (**Fig. 6D**), we separately analyzed cells from the CEPshD lineage (dorsal dataset) and CEPshV lineage (ventral dataset) as two independent datasets (see **Fig. 2A**). For each dataset, two complementary approaches were applied: (1) the FindAllMarkers() function from Seurat (v4) was used to identify differentially expressed genes across Seurat clusters, selecting genes with avg_log_2_FC > 0 and p_val_adj < 0.05; and (2) the graph_test() function from Monocle3(Trapnell et al. 2014; Qiu et al. 2017; Cao et al. 2019) was used to identify genes whose expression varies significantly along pseudotime trajectories, using a cutoff of q_value < 0.05. Only genes identified by both methods were retained as dynamically expressed genes. The resulting gene sets from the dorsal and ventral datasets were then combined to generate a union list of dynamically expressed genes in CEPsh glia lineages.

L2-L3 CEPsh glia enriched genes (**Supplemental Fig. S6F**) were obtained from a previously published RNA sequencing study (Katz et al., 2019)(Katz et al. 2019). To identify adult CEPsh glia enriched genes (**Supplemental Fig. S6G**), single-cell gene expression data was obtained from a recently published atlas (Purice et al., 2025)(Purice et al. 2025). CEPsh glia enriched genes were identified using FindMarkers() function from Seurat (v4) with avg_log2FC > 0 and p_val_adj < 0.05.

### NanoDam profiling

To prepare animals for NanoDam profiling, strains carrying the NanoDam transgene alone or together with an endogenously GFP-tagged CEH-43 were maintained on *dam–/dcm– E. coli* (NEB C2925) for at least three generations without starvation. Nine 55 mm plates densely populated with gravid adults were bleached to collect embryos. The resulting synchronized L1s were placed onto 90 mm NGM plates seeded with *dam–/dcm– E. coli*, grown at 20°C, and harvested at the L4 stage.

Two biological replicates were obtained for each condition and processed in parallel to minimize batch effects. Genomic DNA was extracted using the QIAamp® DNA Micro Kit (Qiagen), following the manufacturer’s protocol for “Isolation of Genomic DNA from Tissues.” Downstream steps followed a previously published DamID protocol(Marshall et al. 2016). Genomic DNA was digested with DpnI (NEB), and DamID adaptors were ligated using T4 DNA ligase (NEB). Ligated DNA was then digested with DpnII (NEB) and purified using homemade Sera-Mag beads prepared from Cytiva Sera-Mag SpeedBeads and 20% (w/v) PEG-8000 following the method of Rohland and Reich(Rohland and Reich 2012). DNA was amplified with Advantage 2 cDNA Polymerase (Clontech) and quality-checked on a 0.8% agarose gel. Amplicons were purified using the QIAquick PCR Purification Kit (Qiagen), quantified with Qubit, and sonicated to ∼300–400 bp fragments using a Bioruptor Plus. DamID adaptors were removed with AlwI (NEB), followed by one round of bead purification. End-repair was performed using an enzyme mix of T4 DNA polymerase, Klenow fragment, and T4 polynucleotide kinase (NEB), followed by 3′ adenylation with Klenow Fragment (3′→5′ exo–) (NEB) and ligation of indexed sequencing adaptors. Libraries were bead purified twice, amplified with NEBNext PCR Master Mix, purified again, checked for quality on a TapeStation, and pooled for Illumina sequencing.

### NanoDam peak analysis

Demultiplexed data were analyzed using a previously published pipeline (GitHub: https://github.com/AHBrand-Lab/NanoDam_analysis)(Tang et al. 2022). CEH-43 NanoDam signal z-score aggregation plots were generated using SeqPlots (v3.0.12)(Stempor and Ahringer 2016). CEH-43 peaks with FDR <10^-25^ were annotated to nearby genes using both bedtools (v.2.28) and HOMER (*annotatePeaks.pl*, v.4.11). Gene lists obtained from both methods were merged to create a unified list of candidate CEH-43 target genes.

### Comparative analysis with mammalian astrocytes

Mammalian astrocyte single-cell RNA sequencing data were obtained from two previous publications(Wang et al. 2025; Frazel et al. 2023), and re-analyzed according to the workflow described in these papers. Human and mouse astrocyte enriched genes were identified using FindAllMarkers() function in Seurat (v4). Ortholog matching between human and *C. elegans* genes was done with OrthoList 2(Kim et al. 2018), and ortholog matching between mouse and human genes was done with Biomart on Ensembl(Dyer et al. 2025). The module scores of the top 50 mammalian radial glia/astrocyte/neuron enriched genes in *C. elegans* CEPsh glia lineages were calculated with the AddModuleScore() function. The output module scores represent the average expression of the input gene set in each cell, normalized against a background of control genes with similar overall expression levels. The scores reflect the relative enrichment of mammalian astrocyte- or neuron-enriched genes in individual *C. elegans* cells.

*p*-values for pairwise module score comparisons between CEPsh lineage subclusters are listed below (Wilcoxon test with Bonferroni correction):

#### 1. Wang et al., 2025 Human

##### Radial glia enriched genes, compared with Converged_CEPsh glia

2.33 × 10^-80^, 2.20 × 10^-55^, 3.80 × 10^-41^, 1.73 × 10^-60^, 1.17 × 10^-120^, 1.09 × 10^-1^, 1.15 × 10^-5^, 8.77 × 10^-1^, 7.63 × 10^-1^, 1.02 × 10^-35^

##### Astrocyte enriched genes, compared with Converged_CEPsh glia

5.27 × 10^-143^, 4.03 × 10^-163^, 1.50 × 10^-86^, 1.92 × 10^-157^, 3.11 × 10^-202^, 3.61 × 10^-18^, 1.50 × 10^-42^, 8.36 × 10^-36^, 2.33 × 10^-6^, 6.55 × 10^-70^

##### Interneuron enriched genes, compared with URAV

1.91 × 10^-68^, 1.26 × 10^-50^, 1.84 × 10^-49^, 3.53 × 10^-8^, 7.60 × 10^-47^, 3.86 × 10^-7^, 2.00 × 10^-3^, 4.44 × 10^-27^, 4.00 × 10^-3^, 1.22 × 10^-23^

#### 2. Frazel et al., 2023 Human

##### Astrocyte (1) enriched genes, compared with Converged_CEPsh glia

7.35 × 10^-132^, 4.50 × 10^-131^, 7.12 × 10^-77^, 7.65 × 10^-129^, 8.30 × 10^-192^, 1.58 × 10^-39^, 9.56 × 10^-55^, 3.64 × 10^-11^, 7.35 × 10^-7^, 2.62 × 10^-89^

##### Astrocyte (2) enriched genes, compared with Converged_CEPsh glia

6.85 × 10^-43^, 2.47 × 10^-49^, 6.29 × 10^-21^, 8.70 × 10^-28^, 1.64 × 10^-29^, 7.20 × 10^-18^, 2.75 × 10^-22^, 2.48 × 10^-11^, 8.53 × 10^-4^, 4.90 × 10^-40^

##### Neuron enriched genes, compared with URAV

2.21 × 10^-4^, 3.42 × 10^-13^, 9.16 × 10^-1^, 6.00 × 10^-3^, 5.23 × 10^-34^, 5.70 × 10^-14^, 4.72 × 10^-7^, 3.20 × 10^-30^, 2.32 × 10^-7^, 1.86 × 10^-1^

#### 3. Frazel et al., 2023 Mouse

##### Astrocyte enriched genes, compared with Converged_CEPsh glia

3.47 × 10^-75^, 1.90 × 10^-69^, 5.04 × 10^-42^, 2.33 × 10^-47^, 6.60 × 10^-118^, 3.50 × 10^-47^, 3.47 × 10^-75^, 1.18 × 10^-55^, 8.42 × 10^-1^, 9.93 × 10^-55^

##### Neuron enriched genes, compared with URAV

2.13 × 10^-31^, 2.98 × 10^-44^, 2.85 × 10^-13^, 2.22 × 10^-32^, 1.20 × 10^-5^, 1.61 × 10^-50^, 9.95 × 10^-48^, 3.80 × 10^-2^, 1.58 × 10^-42^, 7.37 × 10^-25^

Genes enriched in *C. elegans* CEPsh glia and in different human/mouse astrocytes were identified (average log_2_FC >0.5 and adjusted p-value <0.05). Shared orthologs were identified.

### RNAi screen

The RNAi screen was performed using the standard RNAi feeding method(Conte Jr. et al. 2015). RNAi bacterial clones were obtained from the Ahringer RNAi feeding library(Kamath and Ahringer 2003). Overnight cultures of RNAi bacteria were resuspended in M9 buffer and plated onto NGM plates containing 1 mM IPTG (Sigma). Plates were dried overnight. L4-stage *C. elegans* were first transferred to empty plates to remove residual bacteria from their bodies and then moved to the RNAi plates. For each genotype, L1 larvae and L4 larvae were scored for CEPsh glia reporter expression.

### Fluorescence imaging and quantification

Fluorescence imaging was performed using a Zeiss Imager M2 microscope equipped with an AxioCam MRm 1.4 camera and a 63x/1.4 NA oil immersion objective. Confocal images were acquired using a Zeiss LSM 900 inverted laser scanning confocal microscope with a 63x/1.4 NA oil objective. Animals were immobilized in 200 mM sodium azide either on 1% agar pads or with 20 µm polystyrene microbeads. For quantification of *glt-1p::gfp* expression in **Fig. 6Q**, regions of interest were manually drawn around nuclei of CEPsh glia. Mean fluorescence intensity was measured at the central z-plane of the cell body. Fluorescence intensities from wild-type and *ceh-43(ot406)* animals were normalized to the average intensity of the wild-type group.

### iSPIM imaging and lineage tracking

A custom-built scanning iSPIM microscope was used for embryo imaging(Insley and Shaham 2018; Wu et al. 2011). Two-cell stage embryos were placed on a dried poly-L-lysine spot in the center of a Fisherbrand 24 × 50-1 microscope cover glass (Thermo Fisher Scientific, Cat. 12-545-F). Embryos were immersed in M9 buffer within an imaging chamber and imaged for 10 hours at 75-second intervals. Lineage tracking was performed with StarryNite(Bao et al. 2006), and lineages were visualized with AceTree(Boyle et al. 2006).

### CRISPR/Cas9 genome editing

Generation of endogenous *glt-1::SL2::GFP::H2B* reporter and motif mutations was performed using CRISPR/Cas9 using Cas9, tracrRNAs, and crRNAs from IDT, as previously described(Eroglu et al. 2023).

- *ns1117[glt-1::SL2::GFP::H2B]*: crRNA (AATTCGGTGTGATTAAAATC), ssODN (prepared by enzymatic digestion of PCR product into single-stranded DNA as previously described(Eroglu et al. 2023); see **Supplemental Information**).
- *ns1119* [153-bp region in the *glt-1* promoter replaced with an XhoI site]: crRNA #1 (AATTCGGTGTGATTAAAATC), crRNA #2 (AGTACTTTAGATAGACGACA), ssODN (AAAAGTGTCACCTCAACTTTATGTCACATTCTGTACTCGAGACAAGGGAGGGGGAGGGAAGGGGTAAAATACTAGA, ordered as IDT Ultramer).

### CEPsh glia rescue experiment

Rescue experiments in *ceh-43(ot406)* mutants were performed using the following DNA injection mixes:

#### Fosmid-based rescue

A *ceh-43* fosmid (15 ng/µL) was co-injected with *elt-2::mCherry* plasmid (10 ng/µL).

#### Genomic DNA rescue

*ceh-43p::ceh-43* genomic PCR product (amplified using oSL265 and oSL398, 10 ng/µL) was co-injected with pSL037(*nhr-2p::his-24::mCherry*, 15 ng/µL) and *elt-2::mCherry* plasmid (10 ng/µL).

#### *ceh-43* cDNA driven by different promoters

pSL056(*sox-2p::ceh-43 cDNA::SL2::his-24::mCherry*) was co-injected with *elt-2::mCherry* plasmid (10 ng/µL) and pSL046(*nhr-2p::his-24::TagBFP(I174A)*, 10 ng/µL). The *pros-1* promoter (amplified using oSL453 and oSL457 from the genome) was fused to *ceh-43 cDNA::SL2::his-24::mCherry::let-858 3’UTR* by PCR. The fusion product (10 ng/µL) was injected with *elt-2::mCherry* plasmid (10 ng/µL). pSL047(*mir-228p::ceh-43 cDNA::SL2::his-24::mCherry*, 10 ng/µL) was co-injected with *elt-2::mCherry* plasmid (10 ng/µL).

The rescue experiment in *ztf-29(gk561)* mutants was performed using the following DNA injection mixes:

#### Genomic DNA rescue

The *ztf-29* genomic locus (*ztf-29p(3kb)::ztf-29::ztf-29 3’UTR*, cloned into pKB001, 15 ng/µL) was co-injected with *elt-2::mCherry* plasmid (10 ng/µL).

All injection mixes were supplemented with pBluescript plasmid to reach a final DNA concentration of 100 ng/µL.

### Mosaic analysis

We performed mosaic analysis with two different *ceh-43* mutants. *ceh-43(ot406)* mutant animals were generated that harbor extrachromosomal transgenic DNA arrays that are stochastically transmitted to daughter cells during cell divisions. *ceh-43* fosmid was injected (15 ng/µL) together with a ubiquitous mCherry co-injection marker (*nhr-2p::his-24::mCherry*, 10 ng/µL) to mark the existence of the array. In a separate strategy, *mir-228p::ceh-43(cDNA)::SL2::H1-mCherry* (10 ng/µL) was injected, with H1-mCherry indicating sites *mir-228p* expression. For the embryonic lethal *ceh-43(tm480)* mutant, *ceh-43(tm480)/mT1* animals were injected with *ceh-43* fosmid (5 ng/µL) and *nhr-2p::his-24::mCherry* (15 ng/µL). Rescue array positive and mT1 balancer negative animals were picked as *ceh-43(tm480)* homozygous mutant animals for mosaic analysis.

### Heat-shock induction of *ceh-43* expression

A heat-shock construct (*hsp16.41p::ceh-43 cDNA::SL2::his-24::mCherry::let-858 3’UTR*) was injected at 10 ng/μl, together with *elt-2*::mCherry (10 ng/μl) as a co-injection marker. Mixed-stage transgenic animals were heat-shocked at 32 °C for 10 minutes, allowed to recover at 20 °C for 4-6 hours, and then imaged and scored for reporter induction.

### Cell culture

Mouse primary astrocytes (isolated from E14 or E15 mice, Lonza, Cat. No. M-ASM-330) were cultured following a protocol adapted from the manufacturer’s instructions. Briefly, cells were seeded onto multi-well plates pre-coated with poly-D-lysine (Sigma, Cat. No. P1024-50MG) and laminin (Gibco, Cat. No. 23-017-015) immediately after thawing. Cultures were maintained at 37 °C in a humidified incubator with 5% CO_2_. Eight hours after initial attachment, the medium was replaced with fresh growth medium. A half-medium change was performed on day 7, followed by medium changes every 4–5 days until the cultures reached confluence. Cells were then passaged once onto coverslips and maintained for 8 days before fixation to analyze immature astrocytes.

The growth medium was the AGM™ Astrocyte Medium BulletKit (Lonza, Cat. No. CC-3186), which includes fetal bovine serum (FBS), L-glutamine, GA-1000, ascorbic acid, human epidermal growth factor (hEGF), and insulin.

### Immunostaining and confocal imaging

Immunostaining protocol on cultured astrocytes is as follows. Primary astrocytes (isolated from E14 or E15 mice, Lonza, Cat. M-ASM-430) grown on coverslips were fixed with 4% paraformaldehyde for 20 minutes and rinsed three times with DPBS (Gibco, Cat. 14190-144, 10 minutes each). Cells were permeabilized in 0.2% Triton X-100 (Sigma, Cat. T8787) prepared in 10% normal goat serum (NGS, Vector Laboratories, Cat. S-1000-20) for 15 minutes, followed by three DPBS washes (5 minutes each). After blocking with 10% NGS for 2 hours at room temperature, cells were incubated overnight at 4 °C with the following primary antibodies: rabbit anti-DLX1 (1:500; Cell Signaling Technology, Cat. 96585), rabbit anti-DLX2 (1:500, Proteintech, Cat. 26244-1-AP), and chicken anti-GFAP (1:500; Aves Labs, Cat. GFAP). The next day, coverslips were washed three times with DPBS (5 minutes each) and incubated for 1 hour at room temperature with the following secondary antibodies (all at 1:1000): Alexa Fluor™ 488 Goat anti-Rabbit IgG (H+L) (Invitrogen, Cat. A-11008) and Alexa Fluor™ 594 Goat anti-Chicken IgY (H+L) (Invitrogen, Cat. A-11042). Cells were then washed three times with 1× PBST (DPBS containing 0.05% Tween-20 (Sigma, Cat. P9416), 5 minutes each) and incubated with Hoechst 33342 (1:10,000 dilution; Cat. 62249) for 5 minutes at room temperature, followed by three washes with DPBS (5 minutes each). Coverslips were mounted using SlowFade Gold Antifade Mountant (Invitrogen, Cat. S36940). Images were acquired using a Zeiss scanning confocal microscope (LSM900).

The protocol for immunocytochemistry on mouse brain slices was adapted from a published method(Medrihan et al. 2025). Briefly, mice were perfused intracardially with 20 mL of ice-cold 1× PBS, followed by 20 mL of 4% paraformaldehyde (PFA) in 1× PBS. Brains were post-fixed in 4% PFA for 1-3 hours or overnight. Brains were then cryopreserved in 5% sucrose in 1× PBS for 1 hour, followed by 15% and 30% sucrose. Brains were embedded in OCT compound and stored in a -80 °C freezer until sectioning. Cryostat sections (16-20 µm thick) were collected and stored at -80 °C in air-tight boxes until use. For staining, sections were air-dried for 2 hours and washed three times (5 min each) in 1× PBS. Blocking was performed for 1 hour at room temperature in 10% normal goat or donkey serum with 0.2% Triton X-100 in 1× PBS. Primary antibodies, diluted in antibody diluent (5% normal goat serum (Vector Laboratories, Cat. S-1000-20) and 0.2% Triton X-100 in 1× PBS), were applied overnight at 4 °C. After washing the sections in 1× PBS with 0.2% Triton X-100, sections were incubated for 1 h at room temperature with appropriate fluorescent secondary antibodies and nuclei stain DAPI, diluted in antibody diluent, in an airtight black humidified chamber box. Sections were then washed in 1× PBS, coverslipped with anti-fade mounting medium, and imaged using a confocal microscope.

Following antibodies and dilutions were used in this study: DLX1 (Cell Signaling, Cat. 96585, dilution 1:500), DLX2 (Proteintech, Cat. 26244-1-AP; dilution 1:250), GFP (Abcam, Cat. ab13970, 1:500), and secondary antibodies: Alexa Fluor 568 goat-anti-rabbit IgG (H+L) highly cross-absorbed (Invitrogen, Cat. A11036), Alexa Fluor 488 goat anti-chicken IgY (H+L) highly cross-absorbed (Invitrogen, Cat. A32931) diluted at 1:500 in antibody diluent.

Confocal images were acquired using a Zeiss LSM 900 microscope. Maximum intensity projections were generated in Fiji from z-stacks with 0.5-1 µm optical sections. Images were cropped and adjusted for brightness and contrast in Photoshop.

### Quantification and statistical analysis

For quantification of *glt-1p::gfp* expression shown in Figure 6Q, regions of interest (ROIs) were manually drawn around the nuclei of CEPsh glia based on the *mir-228p::2xNLS::iBlueberry* channel. Mean fluorescence intensity was measured at the central z-plane of the cell body. Fluorescence intensities from both wild-type and *ceh-43(ot406)* animals were normalized to the average intensity of the wild-type group.

Statistical analysis was performed with GraphPad Prism 10 (v10.5.0) and R (v4.2.2). All statistical details can be found in figure legends.

### Data availability

Raw single-cell RNA-seq data and NanoDam sequencing data have been deposited to DDBJ at PRJDB37786 and is publicly available as of the date of publication. Processed data have been deposited to Zenodo (DOI: 10.5281/zenodo.17114839) and is publicly available as of the date of publication. Plasmids, strains, and microscopy data generated in this study are available by contacting Shai Shaham.

### Code availability

Original scripts have been deposited to Zenodo (DOI: 10.5281/zenodo.17114839) and are publicly available as of the date of publication.

## Acknowledgements

We thank members of the Shaham lab and Dr. Cori Bargmann for comments on the manuscript and discussions; Dr. John I. Murray for sharing the code for embryo time analysis; Dr. Andrea Brand for sharing the NanoDam plasmid; and Dr. Peter Meister for sharing DamID strain IS3675. Some strains were provided by the CGC, which is funded by NIH Office of Research Infrastructure Programs (P40 OD010440). FACS was performed at the Rockefeller University Flow Cytometry Resource Center (RRID:SCR_017694), and single-cell library preparation and sequencing was performed at the Rockefeller University Genomics Resource Center (RRID:SCR_020986). Some analyses were performed on computational resources from the Rockefeller University High Performance Computing Resource Center (RRID:SCR_025889). This work was funded by NIH grant R35NS105094 to S.S. S.L. is a Kavli Neural Systems Institute graduate fellow.

## Author contributions

S.L. and S.S. conceptualized the study. S.L. performed all scRNA-seq and NanoDam experiments and conducted the computational analyses with conceptual input from S.S. S.L. performed functional experiments. K.B. carried out the *ztf-29* mutant experiment. J.J.T. characterized AIY neuron defects. Y.A.K. performed the DLX1/2 cell culture antibody staining experiments. A.M. performed antibody staining on mouse brain slices. S.L. wrote the original manuscript draft. S.L. and S.S. reviewed and edited the manuscript. S.S. acquired funding and supervised the study.

