## Supplemental Figures for "Radial-glia-to-astrocyte trans-differentiation and astrocyte transcriptional convergence are coordinated by CEH-43/DLX in *C. elegans*"

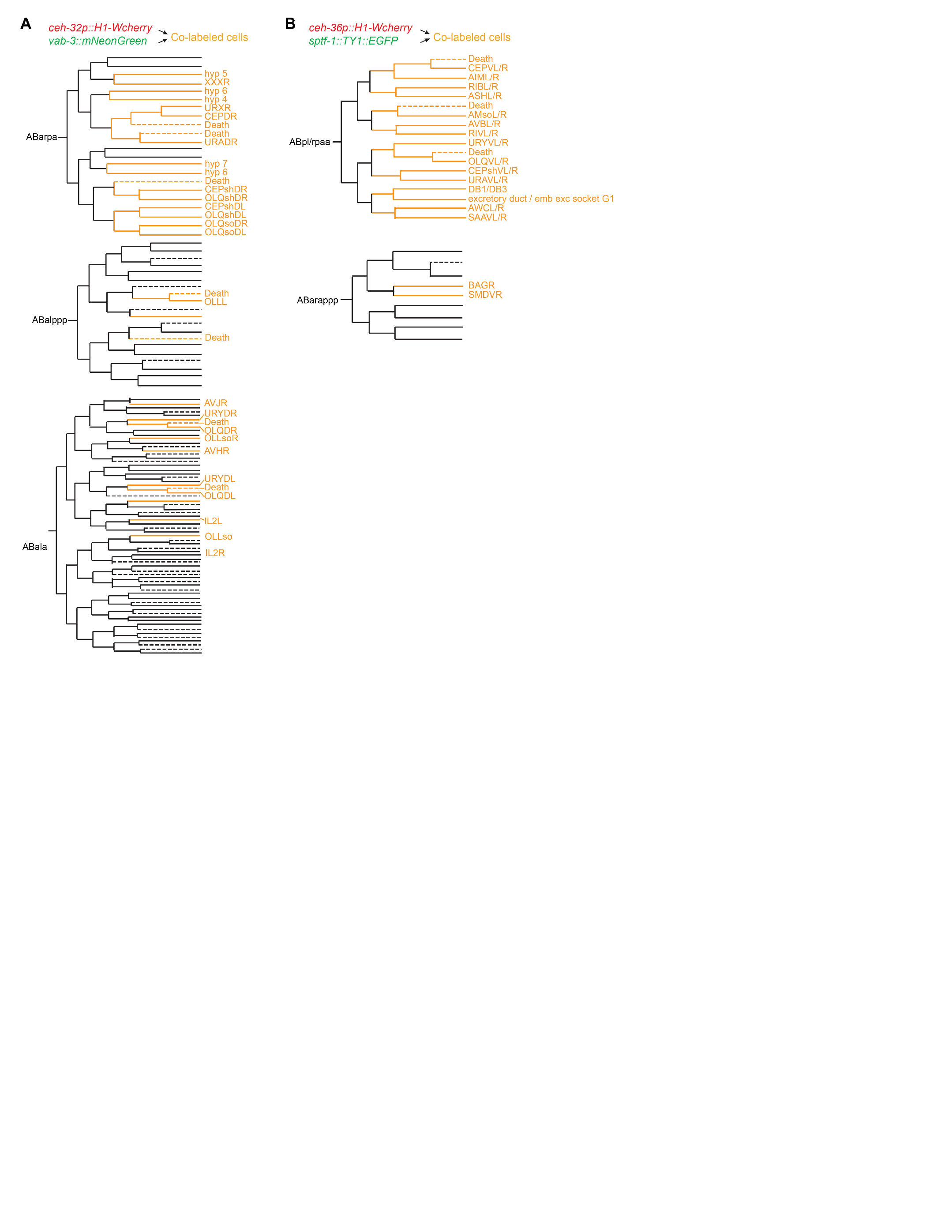

### Supplemental Figure S1. Predicted expression patterns of fluorescent reporter combinations labeling dorsal and ventral CEPsh glia lineages.

**A,B**, Predicted co-labeled cell types for the dorsal (**A**) and ventral (**B**) datasets, based on previously published reporter expression data (Ma et al. 2021).

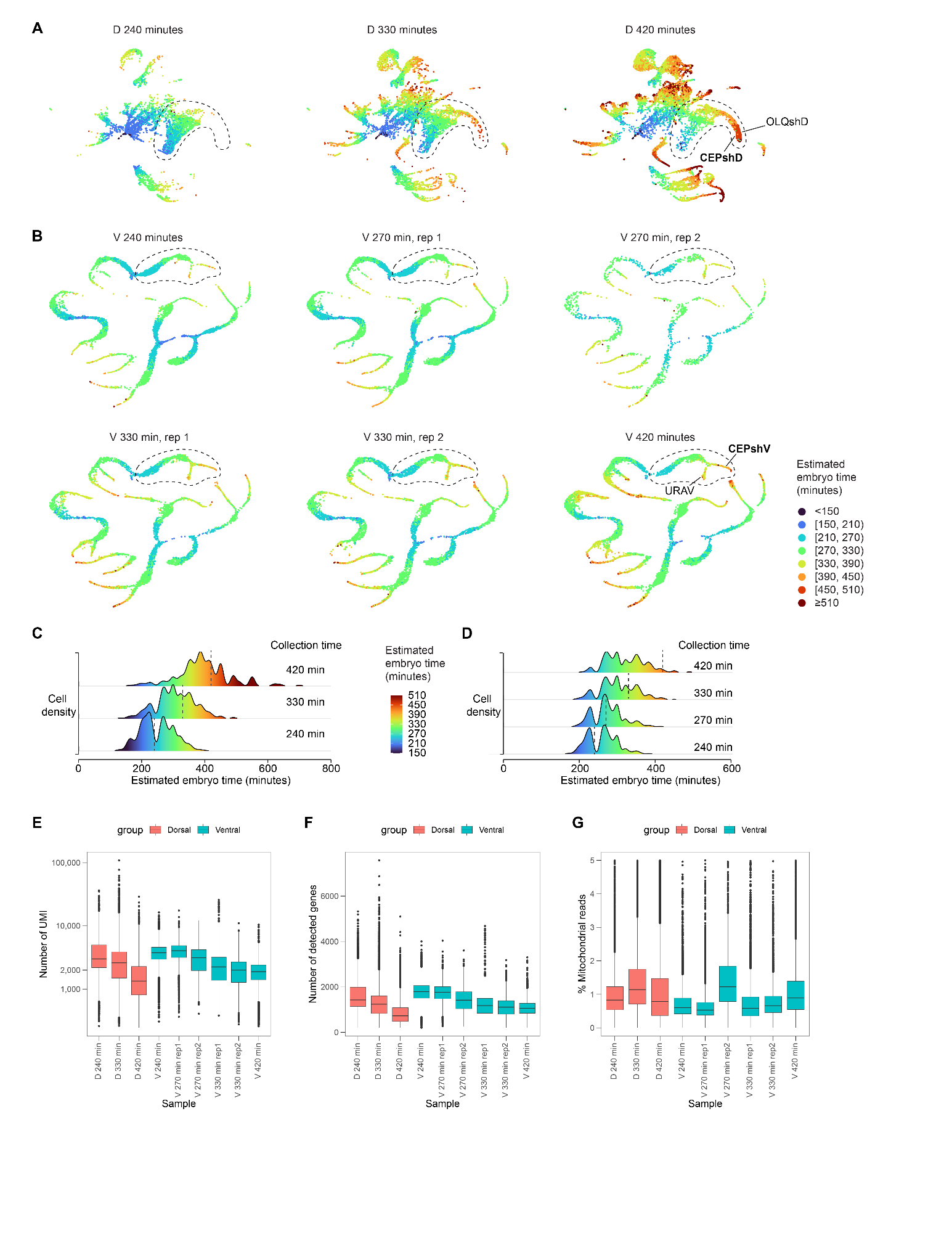

### Supplemental Figure S2. scRNA-seq quality control and developmental timing distribution.

**A,B**, UMAP projections of cells from dorsal (**A**) or ventral (**B**) samples at the indicated time points post-first cell division. Colors indicate estimated embryo time. The sub-lineage containing CEPsh glia, along with their direct sister and progenitor cells, is circled.

**C,D**, Ridge plots of estimated embryo time distributions for cells in the dorsal (**C**) and ventral (**D**) datasets. Dashed vertical lines, sample collection times. For the ventral dataset, embryo collections at 270 and 330 minutes were performed in two independent replicates.

**E-G**, Quality control metrics for scRNA-seq samples: (**E**) number of UMIs per cell, (**F**) number of detected genes per cell, and (**G**) percent mitochondrial reads per cell.

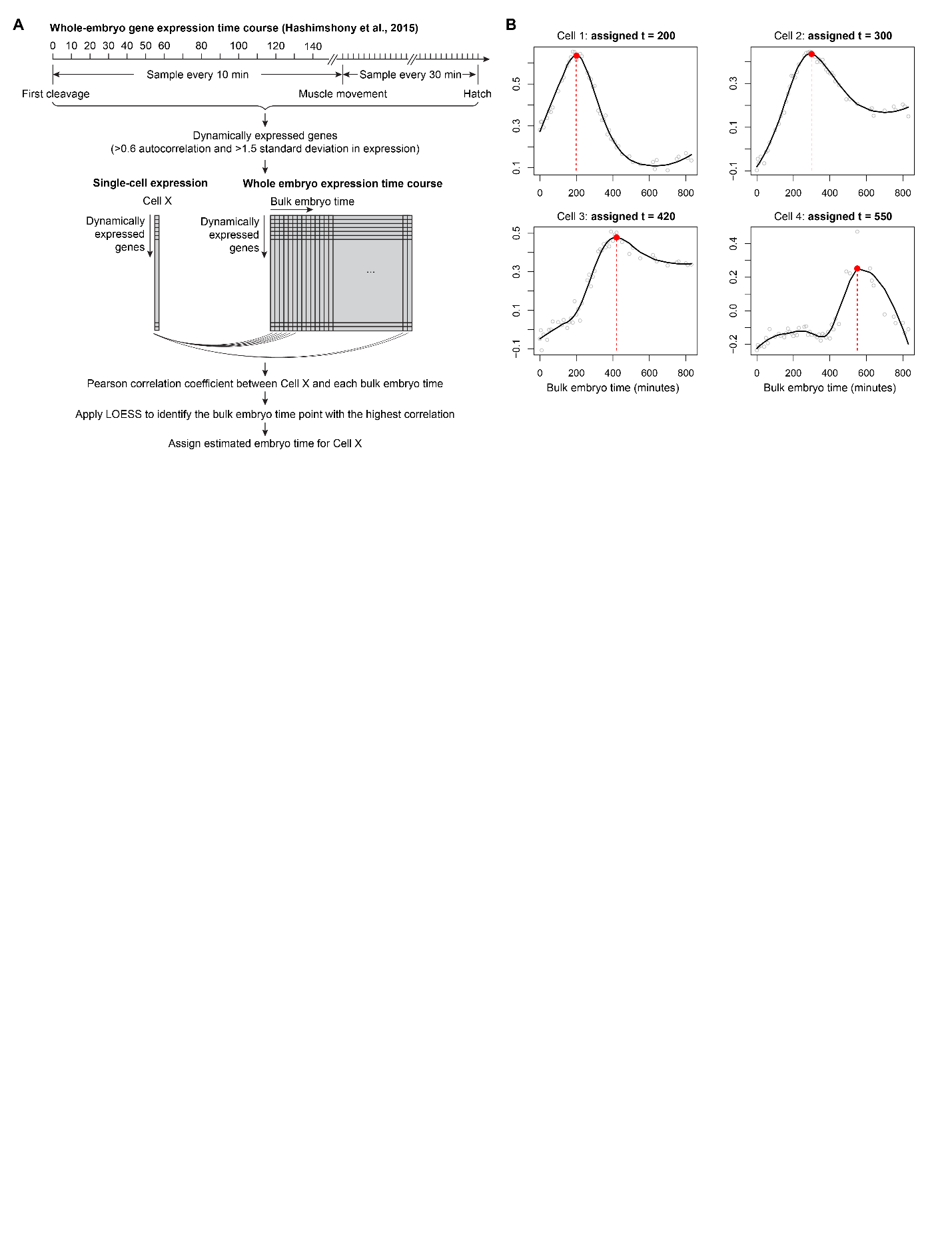

### Supplemental Figure S3. Embryo time estimation.

**A**, Workflow of embryo time estimation. Dynamically expressed genes were identified from a whole-embryo time-course dataset (Hashimshony et al. 2015). For each single cell, expression levels of these genes were correlated with bulk embryo time points using Pearson correlation. After LOESS smoothing, the time point corresponding to the peak of the smoothed correlation curve was assigned as the cell’s estimated embryo time.

**B**, Example embryo time estimation for four single cells. Open circles, Pearson correlation coefficients across time. Black curve, LOESS-smoothed fit. Red circle, the correlation maximum used to define the estimated embryo time.

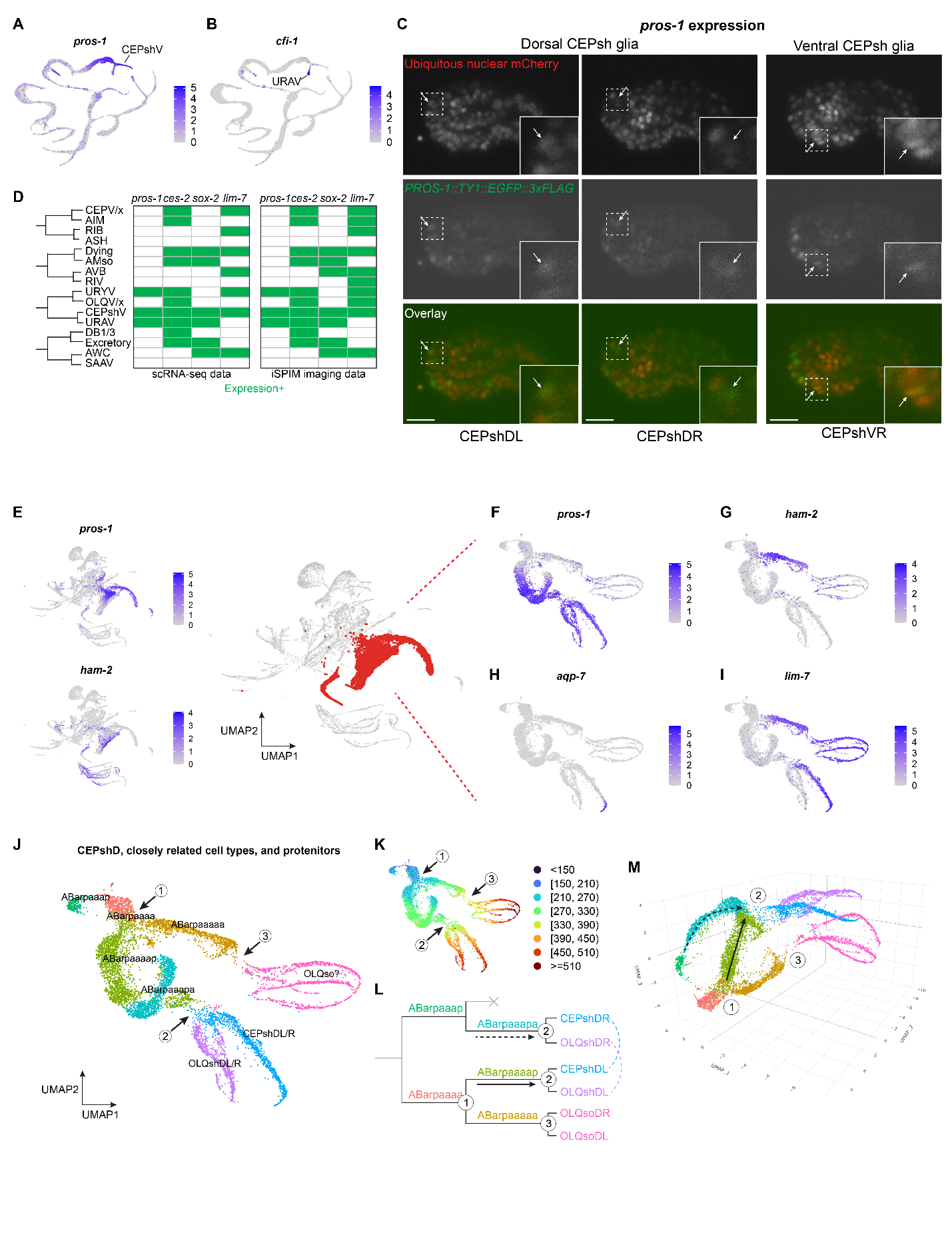

### Supplemental Figure S4. Identification of CEPsh glia lineages and validation of marker gene expression.

**A,B**, Expression patterns of *pros-1* (a CEPshV marker) (**A**), and *cfi-1* (a URAV marker) (**B**).

**C**, iSPIM imaging with lineage tracking showing *PROS-1::TY1::EGFP::3xFLAG* expression in dorsal and ventral CEPsh glia. Insets show enlarged view of the dash line-boxed region highlighting dorsal (CEPshDL, CEPshDR) and ventral (CEPshVR) CEPsh glia (arrows). Scale bar, 10 µm.

**D**, iSPIM imaging validating marker gene expression in lineages related to ventral CEPsh glia. n = 1, 2, 1, 3 embryos for the respective datasets.

**E**, Expression patterns of *pros-1* and *ham-2* in the dorsal dataset, identifying CEPshD glia lineages.

**F-I**, Expression patterns of *pros-1* (**F**), *ham-2* (**G**), *aqp-7* (**H**), *lim-7* (**I**) in CEPshD glia lineages.

**J,K**, UMAP projections showing cell type annotations (**J**) and estimated embryo times (**K**) in CEPshD glia lineages. Cell division events are indicated by numbers (see also (**L**) and (**M**)). The OLQso clusters may represent OLQso glia as well as another socket glial type (CEPso/ILso), which cannot be definitely verified from available gene expression data.

**L**, Lineage diagram of CEPshD glia lineages. Solid and dash arrows indicate two converging CEPsh progenitor lineages (see also panel (**M**)).

**M**, 3D UMAP projection of CEPshD glia lineages. The two CEPshD progenitor lineages (solid and dash lines, respectively) converge and give rise to a shared CEPshD glia cluster.

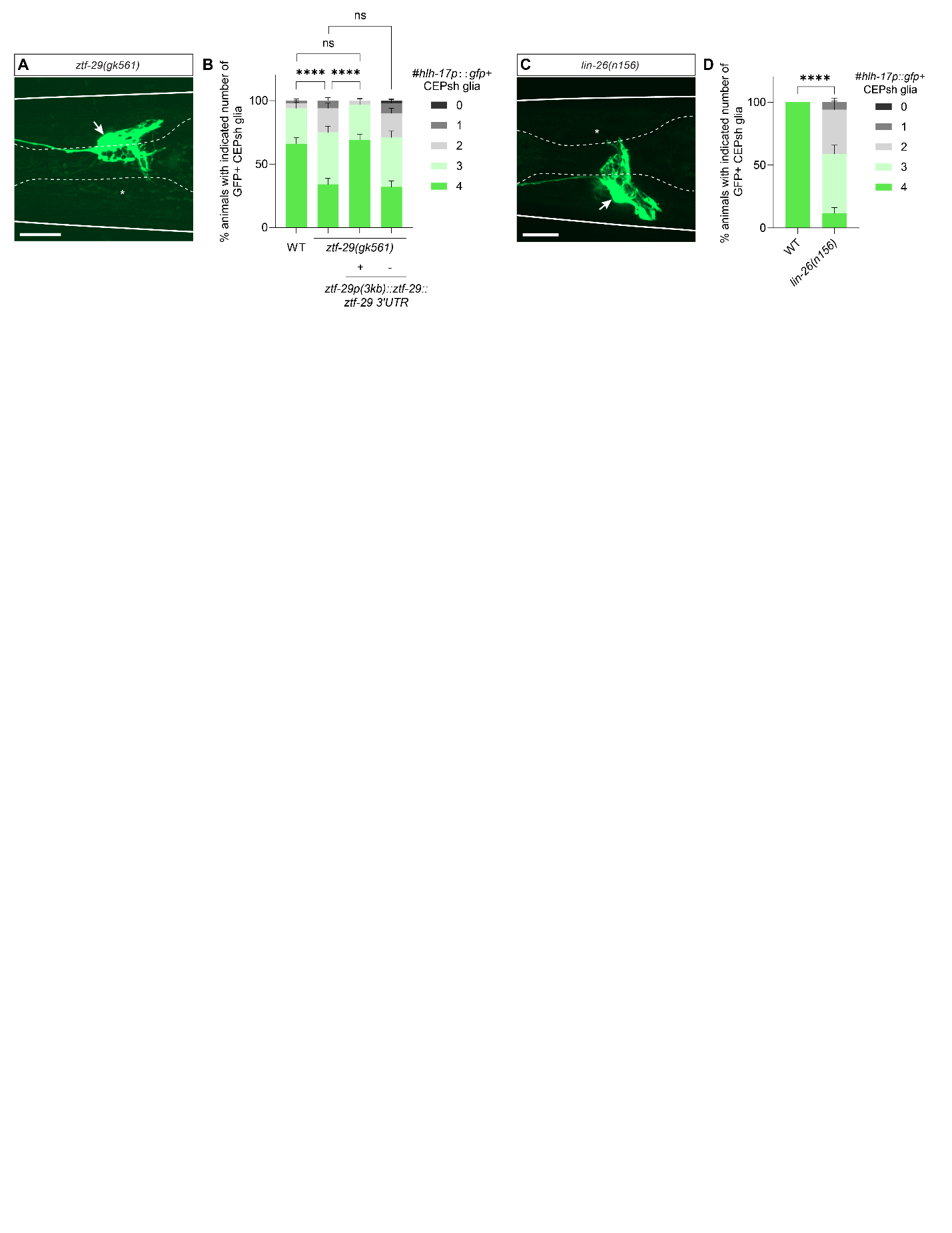

### Supplemental Figure S5. Loss of CEPsh glia reporter expression in lin-26 and ztf-29 mutants.

**A**, Representative confocal image of *hlh-17p::gfp* expression in a *ztf-29(gk561)* mutant L4 larva. Arrow, CEPsh glia. Asterisk, loss of *hlh-17p::gfp* expression in CEPsh glia.

**B**, Quantification of *hlh-17p::gfp* expression in *ztf-29(gk561)* mutants and rescue lines. n = 100 animals per genotype from three independent scoring sessions. *****p* < 0.0001 by Fisher’s exact test.

**C**, Same as in (**A**) but in *lin-26(n156)* mutants.

**D**, Same as in (**B**) but for *lin-26(n156)* mutants.

Scale bars, 10 μm. Error bars, mean ± s.e.m.

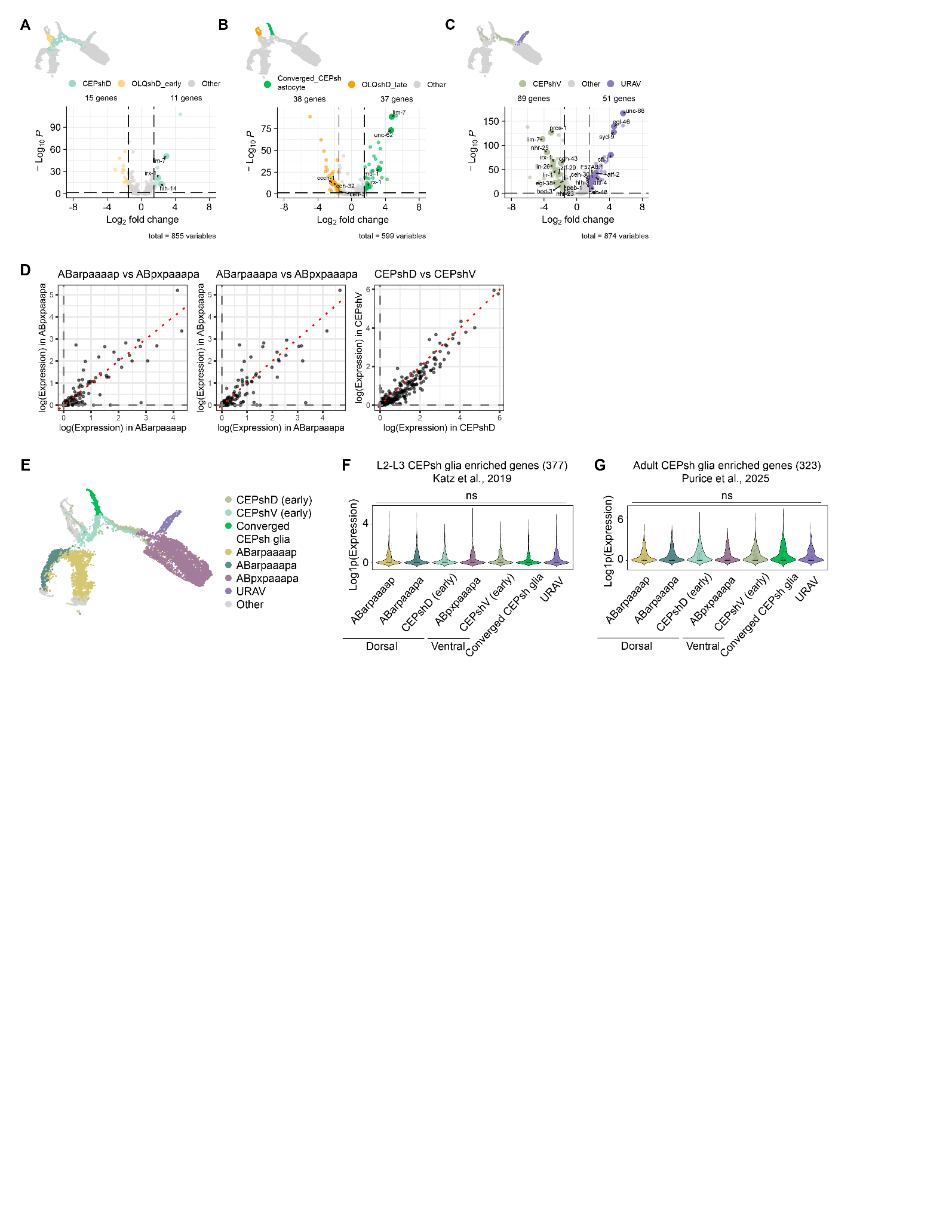

### Supplemental Figure S6. Differential gene expression, converged CEPsh glia-enriched gene expression dynamics, and expression of post-embryonic CEPsh glia-enriched genes in embryonic CEPsh glia lineages.

**A-C**, Differential gene expression analyses showing divergence of CEPsh glia gene expression from their sister cell types. Transcription factors are highlighted. See **Fig. 2C-F** legend for details.

**D**, Scatter plots comparing log-transformed average expression of converged CEPsh glia- or progenitor-enriched genes between dorsal and ventral clusters. Dorsal and ventral CEPsh glia exhibit strong correlation. Red dashed lines, line of equal expression.

**E**, UMAP projection showing cell type clusters from CEPshD and CEPshV lineages analyzed in (**F**) and (**G**).

**F**, Violin plots showing expression of 377 CEPsh-enriched genes identified in L2–L3 larval astrocytes (Katz et al. 2019) across CEPshD and CEPshV lineage clusters. Each dot represents the average expression of one gene. ns, not significant using Kruskal–Wallis test with Bonferroni correction. Note the low or absent expression of these genes in embryonic lineages, suggesting that embryonic and larval CEPsh glia are transcriptionally distinct.

**G**, Same as in (**F**) but for adult CEPsh glia (Purice et al. 2025).

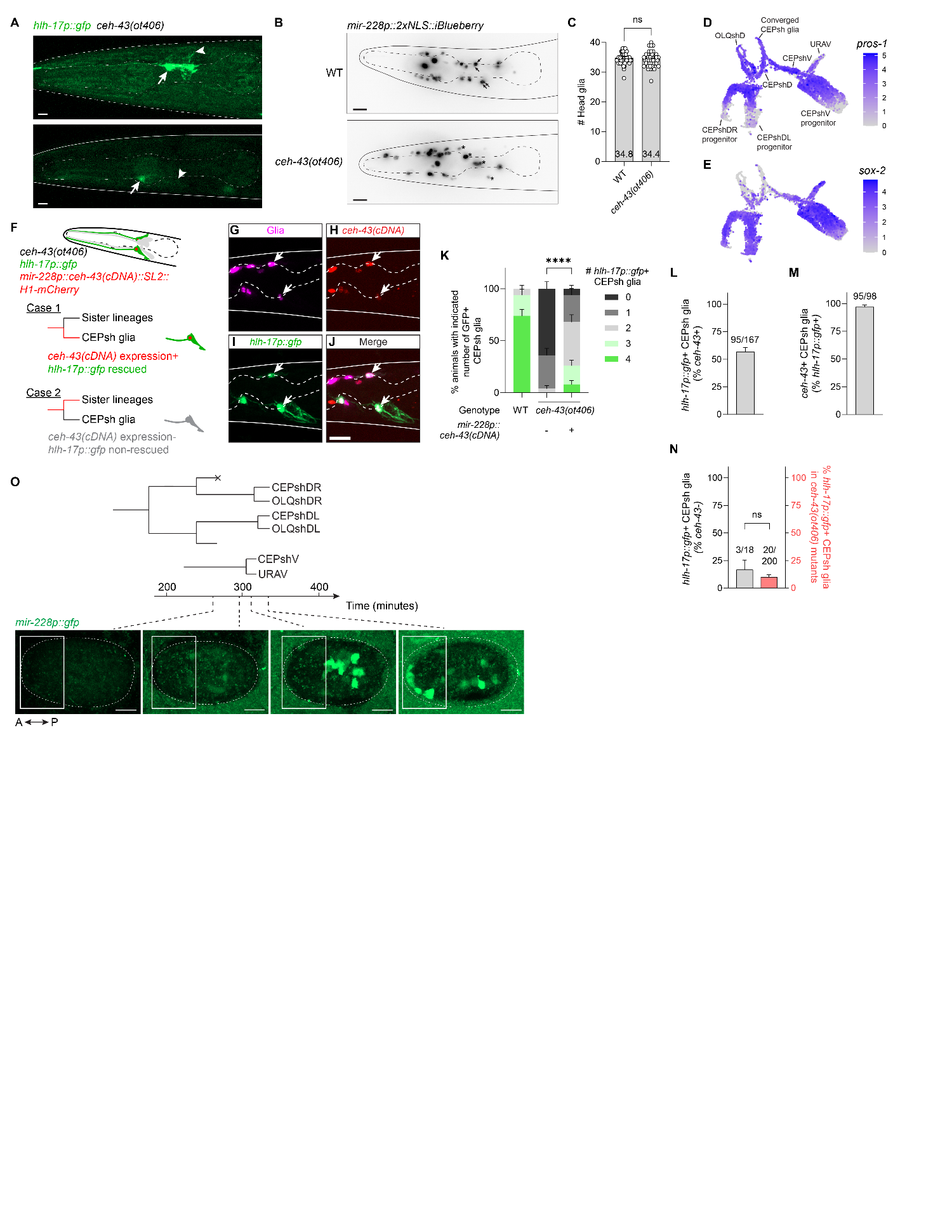

### Supplemental Figure S7. CEH-43 functions cell-autonomously to regulate hlh-17p::gfp expression in CEPsh glia but is not required for their generation.

**A**, Remaining *hlh-17p::gfp*-positive CEPsh glia in two *ceh-43(ot406)* L4 larvae. Arrows, CEPsh glia somata. Arrowheads, CEPsh glia posterior processes.

**B**, Representative maximum-projection confocal images of L4 wild-type and *ceh-43(ot406)* mutant larvae expressing the pan-glial marker *mir-228p::2xNLS::iBlueberry*. Arrows, CEPsh glia in the wild-type larva. Asterisks, putative CEPsh glia in the mutant larva; because CEPsh cell bodies are displaced in the mutant, their identification is uncertain.

**C**, Quantification of total head glia number, showing no significant difference between wild type and *ceh-43(ot406)* mutants. Numbers within columns, average head glia counts (n = 50 for wild-type and 41 for mutant L4 larvae from three imaging sessions). ns, not significant by Mann–Whitney test.

**D,E**, Embryonic expression of *pros-1* (**D**) and *sox-2* (**E**) in CEPsh glia and their progenitors.

**F**, Strategy for mosaic analysis using *mir-228p::ceh-43(cDNA)::SL2::H1-mCherry* rescue construct, which drives pan-glial expression of *ceh-43(cDNA)*.

**G-J**, Maximum-projection confocal image showing CEPsh glia of *ceh-43(ot406)* mutant animals, rescued using a *mir-228p::ceh-43(cDNA)::SL2::H1-mCherry* transgene. Glia are marked with *mir-228p::2xNLS::iBlueberry* (magenta). Arrows, CEPsh glia.

**K**, Pan-glial *ceh-43(cDNA)* expression restores *hlh-17p::gfp* expression in CEPsh glia. n = 50 animals per group from 2-5 independent scoring sessions. *****p* < 0.0001 by Fisher’s exact test.

**L**, Rescue efficiency of *mir-228p::ceh-43(cDNA)* in CEPsh glia.

**M**, Nearly all CEPsh glia expressing *hlh-17p::gfp* also carry the *mir-228p::ceh-43(cDNA)::SL2::H1-mCherry* rescue array.

**N**, Among CEPsh glia lacking *ceh-43* rescue array expression, only 3 of 18 express *hlh-17p::gfp*, not significantly different from non-transgenic *ceh-43(ot406)* mutants. ns, not significant using Fisher’s exact test.

**O**, Expression of the pan-glial reporter *mir-228p::gfp* in developing embryos. Boxed regions highlight the anterior region containing CEPsh progenitors and differentiated CEPsh glia. *mir-228p::gfp* expression is absent in the anterior region during CEPsh progenitor stages and becomes detectable only after CEPsh glia are born.

Scale bar, 10 µm. Error bars, mean ± s.e.m.

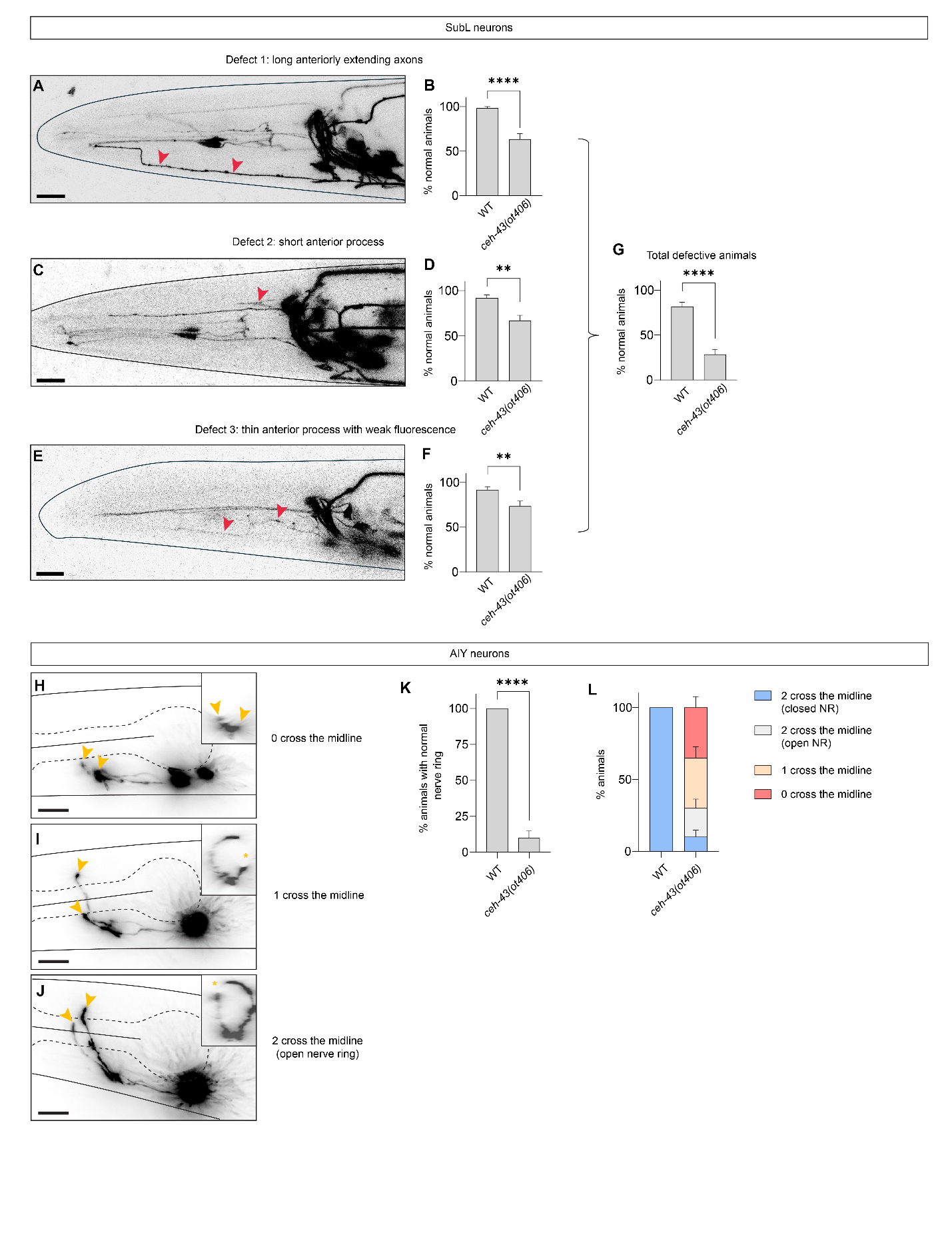

### Supplemental Figure S8. SubL and AIY neuron defects in ceh-43(ot406) mutants.

**A,C,E**, Representative maximum-projection confocal images of *ceh-43(ot406)* mutant animals showing distinct defects in sublateral (SubL) neuron (see wild type in **Fig. 4A**).

**B,D,F,G**, Quantification of SubL neuron defects. n = 60 animals per genotype from three independent imaging sessions. All quantifications were performed on the same cohort of 60 wild-type and 60 *ceh-43(ot406)* mutant animals. ***p* < 0.01, *****p* < 0.0001 by Fisher’s exact test.

**H-J**, Representative maximum-projection confocal images illustrating the three categories of AIY axon extension defects (see wild type in **Fig. 4D**). Arrowheads, short AIY axons; asterisks, gaps in the AIY axon ring.

**K,L**, Quantification of overall AIY axon extension defects. n = 40 animals per genotype from three imaging sessions. *****p* < 0.0001 by Fisher’s exact test.

Scale bar, 10 µm. Error bars, mean ± s.e.m.

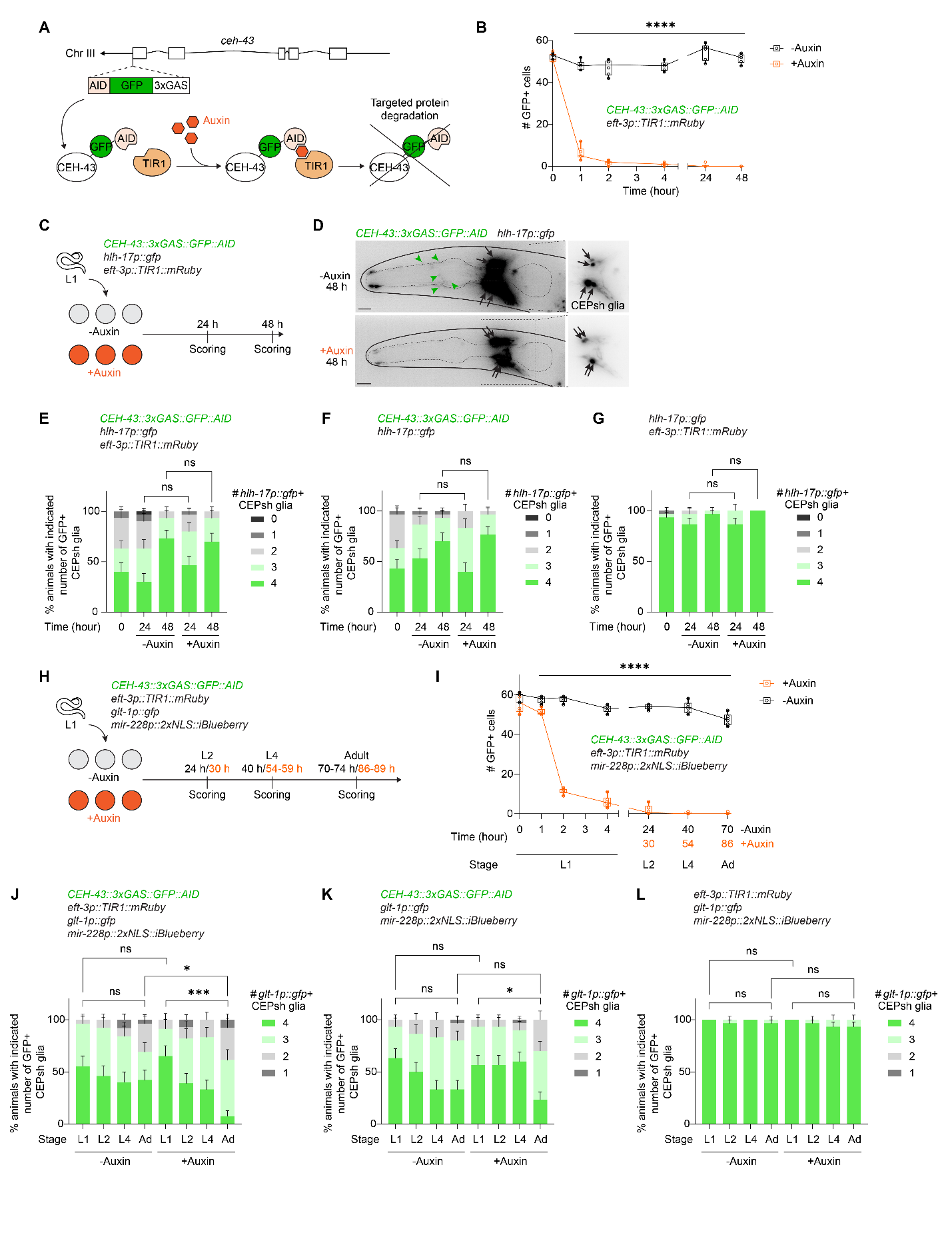

### Supplemental Figure S9. CEH-43 is not required to maintain post-embryonic hlh-17p::gfp or glt-1p::gfp expression.

**A**, Strategy for auxin-inducible CEH-43 protein degradation.

**B**, Time course of CEH-43 protein degradation. n = 6 animals per time point and condition from two duplicate plates in one experiment. Adjusted *p*-values were calculated by Šidák’s multiple comparisons test following two-way ANOVA. *****p* < 0.0001.

**C**, Workflow for testing *hlh-17p::gfp* expression following post-embryonic CEH-43 degradation.

**D**, Representative images of animals grown without (-Auxin) or with (+Auxin) 4 mM Auxin for 48 hours. Boxed areas are enlarged with adjusted contrast to visualize CEPsh glia. Green arrowheads, nuclear CEH-43::GFP signal, which disappears after auxin treatment; black arrows, *hlh-17p::gfp*-positive CEPsh glia. Scale bar, 10 µm.

**E-G**, Quantification of *hlh-17p::gfp*-positive CEPsh glia after auxin treatment in TIR1 + AID (**E**), TIR-only (**F**), and AID-only (**G**) control strains. n = 30 animals from three duplicate plates in one experiment. ns, not significant by Fisher’s exact test.

**H**, Workflow for assessing *glt-1p::gfp* expression following post-embryonic CEH-43 degradation.

**I**, Time course of CEH-43 protein degradation, as in (**B**). n = 6 animals per time point and condition from two duplicate plates in one experiment. Adjusted *p*-values were calculated by Šidák’s multiple comparisons test following two-way ANOVA. *****p* < 0.0001.

**J-L**, Quantification of the number of *glt-1p::gfp*-positive CEPsh glia after auxin treatment in TIR1 + AID (**J**), TIR-only (**K**), and AID-only (**L**) strains. All results are from three duplicate plates in one experiment. Fisher’s exact test, **p* < 0.05, ****p* < 0.001, ns, not significant.

(**J**) n = 27, 26, 25, 26, 23, 28, 24, 26 animals.

(**K**) n = 30 animals per group.

(**L**) n = 27, 30, 30, 30, 26, 30, 30, 30 animals.

Scale bar, 10 µm. Error bars, mean ± s.e.m.

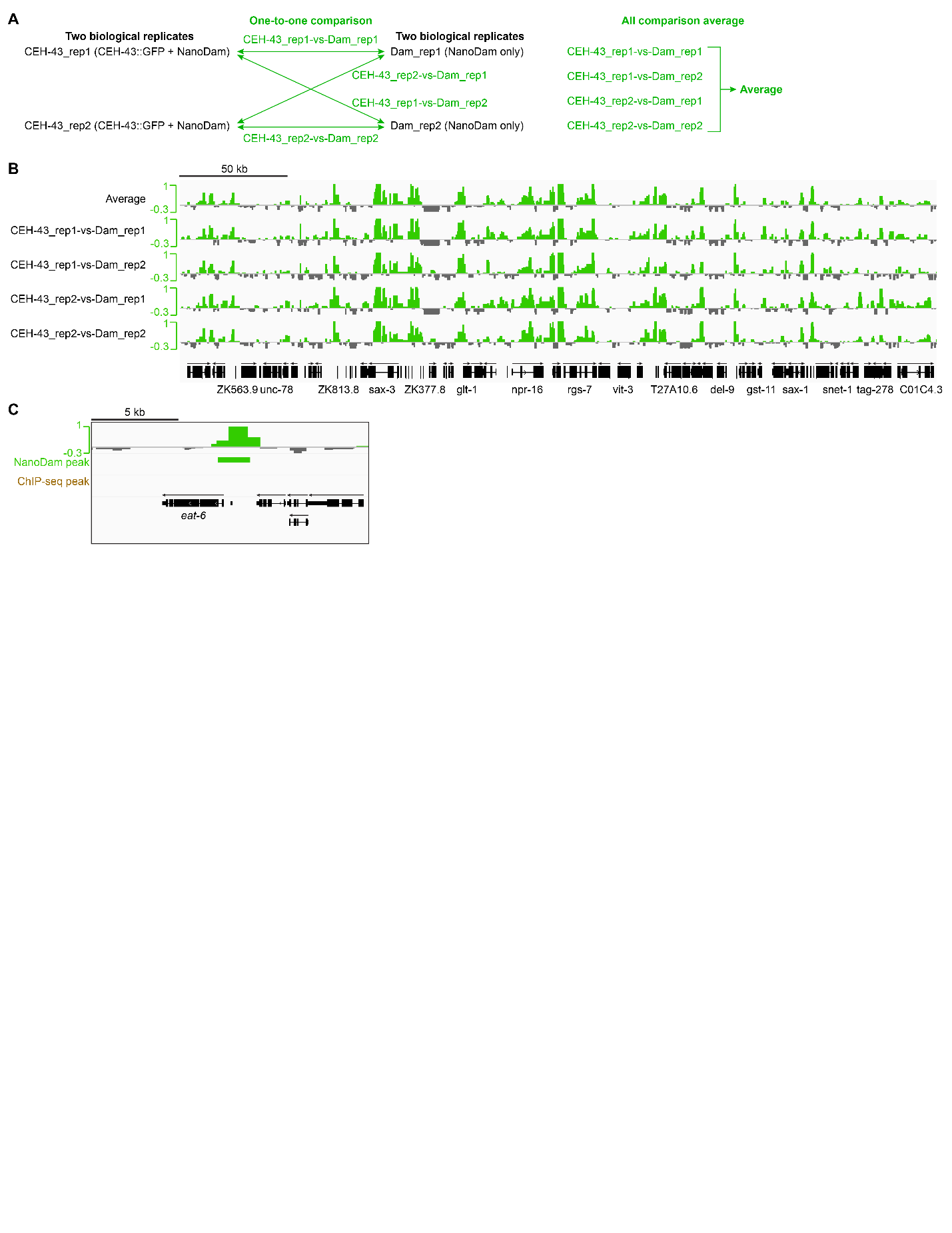

### Supplemental Figure S10. Comparison of CEH-43 NanoDam biological replicates.

**A**, Design of one-to-one comparisons between duplicate CEH-43 NanoDam samples and duplicate NanoDam-only controls. An averaged signal track was generated by combining all one-to-one comparisons.

**B**, Peak tracks from all one-to-one comparisons and the averaged track show strong concordance. A representative genomic region from chromosome X is shown.

**C**, CEH-43 NanoDam binding intensities upstream of *eat-6* coding region.

Arrows in (**B,C**) indicate the direction of coding sequences.

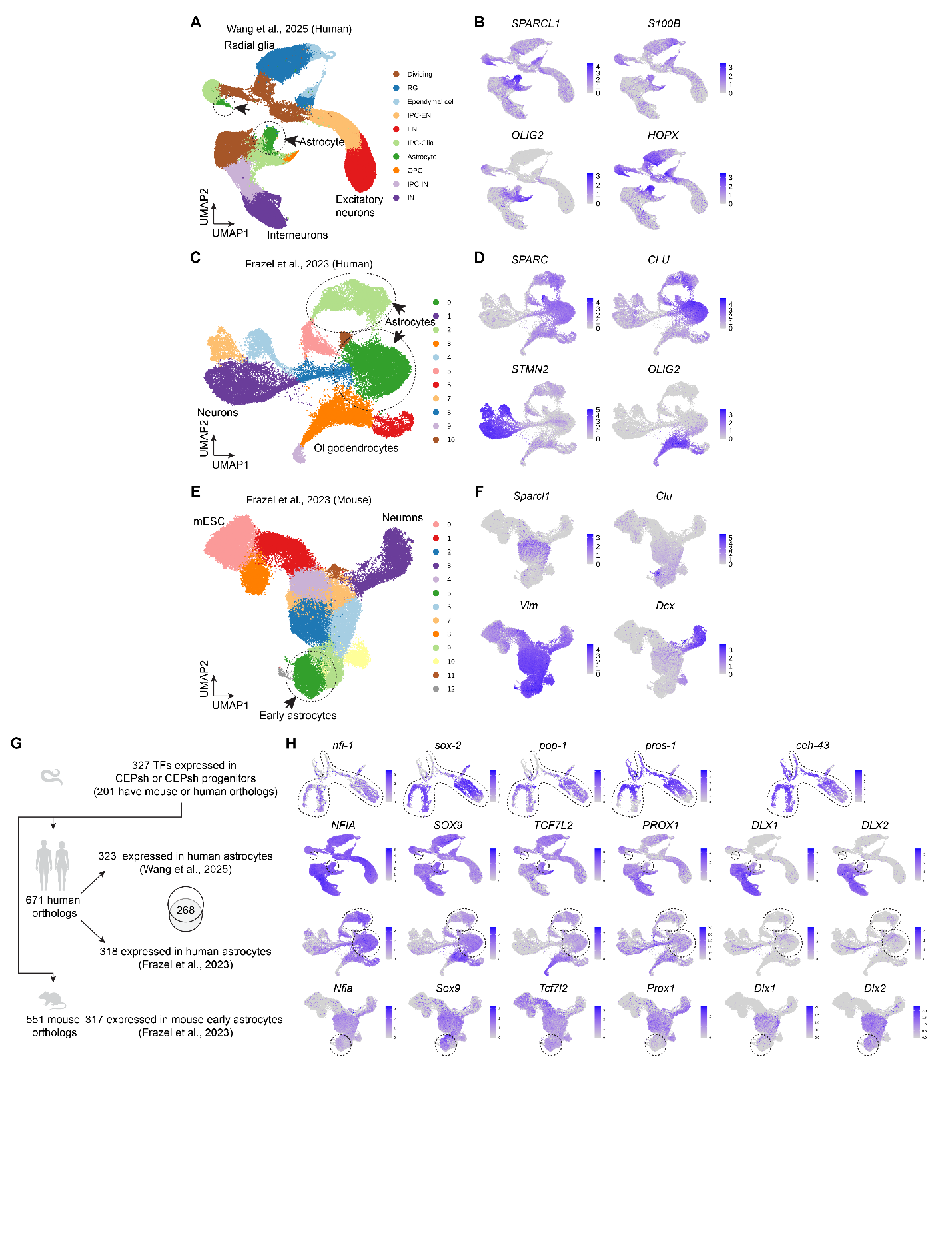

### Supplemental Figure S11. Shared transcription factor expression between C. elegans CEPsh glia and mammalian developing astrocytes.

**A,B**, Reproduced from human cortical glial progenitor *in vitro* differentiation (Extended Data Fig.15 (Wang et al. 2025)). UMAP projections showing original cell-type annotations (**A**) and representative marker gene expression (**B**).

**C,D**, Reproduced from human iPSC-derived glial differentiation scRNA-seq data from (Frazel et al. 2023). UMAP projections of identified clusters (**C**) and representative marker gene expression (**D**).

**E,F**, Reproduced from mouse mESC-derived glial differentiation scRNA-seq data from (Frazel et al. 2023). UMAP projections of identified clusters (**E**) and representative marker gene expression (**F**).

**G**, Overlap between transcription factors expressed in *C. elegans* CEPsh glia or their progenitors and those expressed in mammalian astrocytes.

**H**, Expression patterns of selected transcription factors in CEPsh glia and progenitors, alongside expression of their human and mouse homologs in developing astrocytes.

EPIC database: Expression Patterns In Caenorhabditis. https://epic.gs.washington.edu/Epic2/.
