## Supplemental Information for "Radial-glia-to-astrocyte trans-differentiation and astrocyte transcriptional convergence are coordinated by CEH-43/DLX in *C. elegans*"

ssODN for generating *ns1117[glt-1::SL2::GFP::H2B]*:

GCCTACCAACGGATGACGAGAAGCACACTCATTGAgctgtctcatcctactttcacctagttaactgcttgtcttaaaatctatgcttctctttagtatctaaaattttcctagaagcttacaagtatataaatggtctcttctcaataaaggttgtatatttattcatcttattgaatctgccatttcctcgtttttgcgagtttatataccttccaattttctttctattgtattttcaacttctaattttaattcagggaaactgcttcaacgcatcATGAGTAAAGGAGAAGAACTTTTCACTGGAGTTGTCCCAATTCTTGTTGAATTAGATGGTGATGTTAATGGGCACAAATTTTCTGTCAGTGGAGAGGGTGAAGGTGATGCAACATACGGAAAACTTACCCTTAAATTTATTTGCACTACTGGAAAACTACCTGTTCCATGGgtaagtttaaacatatatatactaactaaccctgattatttaaattttcagCCAACACTTGTCACTACTTTCTgTTATGGTGTTCAATGCTTcTCgAGATACCCAGATCATATGAAACgGCATGACTTTTTCAAGAGTGCCATGCCCGAAGGTTATGTACAGGAAAGAACTATATTTTTCAAAGATGACGGGAACTACAAGACACgtaagtttaaacagttcggtactaactaaccatacatatttaaattttcagGTGCTGAAGTCAAGTTTGAAGGTGATACCCTTGTTAATAGAATCGAGTTAAAAGGTATTGATTTTAAAGAAGATGGAAACATTCTTGGACACAAATTGGAATACAACTATAACTCACACAATGTATACATCATGGCAGACAAACAAAAGAATGGAATCAAAGTTgtaagtttaaacatgattttactaactaactaatctgatttaaattttcagAACTTCAAAATTAGACACAACATTGAAGATGGAAGCGTTCAACTAGCAGACCATTATCAACAAAATACTCCAATTGGCGATGGCCCTGTCCTTTTACCAGACAACCATTACCTGTCCACACAATCTGCCCTTTCGAAAGATCCCAACGAAAAGAGAGACCACATGGTCCTTCTTGAGTTTGTAACAGCTGCTGGGATTACACATGGCATGGATGAACTATACAAACCACCAAAGCCATCTGCCAAGGGAGCCAAGAAGGCCGCCAAGACCGTTACGAAGCCAAAGGACGGAAAGAAGAGACGTCATGCCCGTAAGGAATCATACTCCGTCTACATCTACCGTGTCCTCAAGCAAGTTCATCCAGACACTGGAGTTTCCTCCAAAGCCATGTCTATCATGAACTCTTTTGTCAACGATGTCTTCGAGCGTATTGCTGCTGAAGCATCCCGTCTTGCTCACTACAACAAGCGTTCCACAATCTCATCCCGCGAAATTCAGACCGCTGTCCGTCTGATCCTTCCAGGAGAGCTTGCCAAGCACGCCGTGTCTGAGGGAACCAAGGCCGTTACCAAGTACACTTCCAGCAAGtaatcattgccccagattttaatcacaccgaattatgatttctaaattttct

| **Key Materials and Reagents** | | |
| --- | --- | --- |
| **Reagent or Resource** | **Source** | **Identifier** |
| **Antibodies** |  |  |
| DLX1 (E4T1L) Rabbit mAb | Cell Signaling Technology | 96585 |
| DLX2 Polyclonal antibody (rabbit) | Proteintech | 26244-1-AP |
| Goat anti-Rabbit IgG (H+L) Secondary Antibody, Alexa Fluor™ 488 | Invitrogen | A-11008 |
| Goat anti-Chicken IgY (H+L) Cross-Adsorbed Secondary Antibody, Alexa Fluor™ Plus 488 | Invitrogen | A32931 |
| Goat anti-Rabbit IgG (H+L) Highly Cross-Adsorbed Secondary Antibody, Alexa Fluor™ 568 | Invitrogen | A-11036 |
| Goat anti-Chicken IgY (H+L) Secondary Antibody, Alexa Fluor™ 594 | Invitrogen | A-11042 |
| Anti-Glial Fibrillary Acidic Protein Antibody | Aves Labs | GFAP |
| Anti-GFP antibody | Abcam | ab13970 |
| **Bacterial and virus strains** |  |  |
| *E. coli* OP50 | CGC | https://cgc.umn.edu/strain/OP50 |
| *E. coli dam–/dcm–* | NEB | C2925 |
| **Biological samples** |  |  |
| Mouse CD1 Brain Mixed Astrocytes | Lonza | M-ASM-430 |
| **Chemicals, peptides, and recombinant proteins** |  |  |
| DpnI | New England Biolabs | R0176S |
| DpnII | New England Biolabs | R0543S |
| AlwI | New England Biolabs | R0513S |
| T4 DNA Ligase | New England Biolabs | M0202S |
| T4 DNA polymerase | New England Biolabs | M0203S |
| Klenow fragment | New England Biolabs | M0210S |
| T4 Polynucleotide Kinase | New England Biolabs | M0201S |
| Klenow Fragment (3’→5’ exo-) | New England Biolabs | M0212S |
| EDTA (0.5 M), pH 8.0, RNase-free | Invitrogen | AM9260G |
| NEBNext® High-Fidelity 2X PCR Master Mix | New England Biolabs | M0541S |
| IPTG | Sigma-Aldrich | I6758 |
| Sodium hypochlorite solution | Sigma-Aldrich | 239305-500ML |
| Sucrose (Crystalline/Certified ACS) | Fisher Chemical | S5-3 |
| Chitinase from *Trichoderma viride* | Sigma-Aldrich | C8241 |
| Protease from *Streptomyces griseus* | Sigma-Aldrich | P8811 |
| Bovine Serum Albumin | Sigma-Aldrich | A9647-50G |
| DAPI (4’ ,6-diamidino-2-phenylindole) | Thermo Fisher Scientific | 62248 |
| Poly-L-Lysine, 0.01% solution | Sigma-Aldrich | P4832 |
| Poly-D-lysine | Sigma-Aldrich | P1024-50MG |
| Laminin | Gibco | 23-017-015 |
| DPBS | Gibco | 14190-144 |
| Triton X-100 | Sigma-Aldrich | T8787 |
| Normal Goat Serum Blocking Solution | Vector Laboratories | S-1000-20 |
| Tween-20 | Sigma-Aldrich | P9416 |
| SlowFade™ Gold Antifade Mountant | Invitrogen | S36940 |
| Hoechst 33342 | Thermo Fisher Scientific | 62249 |
| Alt-R™ S.p. HiFi Cas9 Nuclease V3, 100 µg | IDT | 1081060 |
| **Critical commercial assays** |  |  |
| QIAamp® DNA Micro Kit | Qiagen | 56304 |
| QIAquick PCR Purification Kit | Qiagen | 28104 |
| Sera-Mag Speed Bead Magnetic Carboxylate | Cytiva | 6.51521E+13 |
| Advantage 2 cDNA polymerase | Clontech | 639201 |
| Qubit™ 1X dsDNA High Sensitivity | Invitrogen | Q33230 |
| Millex™-SV 5.0 μm Filter Unit (Sterile) | Sigma-Aldrich | SLSV025LS |
| Fisherbrand 24×50-1 microscope cover glass | Thermo Fisher Scientific | 12545F |
| Posi-Click Microcentrifuge Tubes | Denville Scientific Inc. | C2170 |
| AGM^TM^ Astrocyte Growth Medium BulletKit | Lonza | CC-3186 |
| **Cell lines** |  |  |
| P19 cell line | ATCC | CRL-1825 |
| **Oligonucleotides** |  |  |
| *pros-1* promoter forward primer:  GTTTGGTTACGGGATGGC | Sigma-Aldrich | NA |
| *pros-1* promoter reverse primer:  TGATGACGTCACTAGCTA | Sigma-Aldrich | NA |
| *sox-2* promoter forward primer:  CGGAAACGAGTAAAACCC | Sigma-Aldrich | NA |
| *sox-2* promoter reverse primer:  CTTGTTAAACAACAATGA | Sigma-Aldrich | NA |
| *mir-228* promoter forward primer:  GTGACGTCATACTCTTGC | Sigma-Aldrich | NA |
| *mir-228* promoter reverse primer:  AGTTTTTGGGAGGCGACG | Sigma-Aldrich | NA |
| *hsp-16.41* promoter forward primer:  ACGTTGAGCTGGACGGA | Sigma-Aldrich | NA |
| *hsp-16.41* promoter reverse primer:  CTTTCGAAGTTTTTTA | Sigma-Aldrich | NA |
| *ceh-43* genomic locus PCR fragment (*ceh-43p::ceh-43*) forward primer: TTGATGAACTCCTGAAACC | Sigma-Aldrich | NA |
| *ceh-43* genomic locus PCR fragment (*ceh-43p::ceh-43*) reverse primer: AGACATGAGGAAGGGAAAG | Sigma-Aldrich | NA |
| tracrRNA | IDT | 1072532 |
| crRNA for generating *ns1117[glt-1::SL2::GFP::H2B]*: AATTCGGTGTGATTAAAATC | IDT | NA |
| crRNA (#1) for generating *ns1119*: AATTCGGTGTGATTAAAATC | IDT | NA |
| crRNA (#2) for generating *ns1119*: AGTACTTTAGATAGACGACA | IDT | NA |
| **Recombinant DNA** |  |  |
| pSL037 (*nhr-2p::his-24::mCherry::let-858 3'UTR*) | This study, cloned from sequence amplified from *ujIs113* (Murray lab^151^) | NA |
| pSL046 (*nhr-2p::his-24:TagBFP(I174A)::let-858 3'UTR*) | This study | NA |
| pSL047 (*mir-228p::ceh-43 cDNA::SL2::his-24::mCherry::let-858 3'UTR*) | This study | NA |
| pSL048 (*hsp16.41p::ceh-43 cDNA::SL2::his-24::mCherry::let-858 3'UTR*) | This study | NA |
| pSL056 (*sox-2pS::ceh-43 cDNA::SL2::his-24::mCherry::let-858 3'UTR*) | This study | NA |
| pKB001 (*ztf-29p::ztf-29::ztf-29 3’UTR*) | This study | NA |
| WRM0633Cc03 fosmid | *C. elegans* Vancouver fosmid library, Source BioScience | NA |
| **Software and algorithms** |  |  |
| Cell Ranger v7.1.0 | 10x Genomics | <https://www.10xgenomics.com/support/software/cell-ranger/latest> |
| CellBender v0.2.1 | Broad Institute | <https://github.com/broadinstitute/CellBender> |
| Seurat v4 | Integrated analysis of multimodal single-cell data^62^ | <https://satijalab.org/seurat/> |
| SeqPlots v3.0.12 | SeqPlots – Interactive software for exploratory data analyses, pattern discovery and visualization in genomics^140^ | <https://przemol.github.io/seqplots/> |
| StarryNite | Automated cell lineage tracing in *Caenorhabditis elegans*^145^ | <https://github.com/zhirongbaolab/StarryNite> |
| AceTree | AceTree: a tool for visual analysis of *Caenorhabditis elegans* embryogenesis | <https://github.com/zhirongbaolab/AceTree> |

**Mouse and *C. elegans* strains used**

| **Name** | **Genotype** | **Source** |
| --- | --- | --- |
| # 030247 | Mouse: *B6;FVB-Tg(Aldh1l1-EGFP/Rpl10a)JD130Htz/J* | The Jackson Laboratory |
| WormBase: N2; WormBase: WBStrain00000001 | *C. elegans:* Strain N2 | Caenorhabditis Genetics Center |
| RW10584 | *unc-119(ed3) III; zuIs178* *[his-72(1kb 5' UTR)::his-72::SRPVAT::GFP::his-72 (1KB 3' UTR) + 5.7 kb XbaI - HindIII unc-119(+)]; stIs10462[[ceh-32::H1-wCherry::his-24::mCherry + unc-119(+)]* | CGC, Waterston lab |
| SYS633 | *ujIs113 [pie-1p::mCherry::H2B::pie-1 3'UTR + nhr-2p::mCherry::his-24::let-858 3’UTR + unc-119(+)] II; vab-3(dev190([vab-3::mNeonGreen]) X* | CGC(Ma et al. 2021) |
| OS14872 | *vab-3(dev190([vab-3::mNeonGreen]) X; stIs10462 [ceh-32::H1-Wcherry::his-24::mCherry + unc-119(+)]* | This study, generated from strains from Waterston lab and Ma et al., Du lab(Ma et al. 2021) |
| RW10598 | *unc-119(ed3) III; zuIs178 [his-72(1kb 5' UTR)::his-72::SRPVAT::GFP::his-72 (1KB 3' UTR) + 5.7 kb XbaI - HindIII unc-119(+)]; stIs10501 [ceh-36::H1-wCherry::his-24::mCherry + unc-119(+)]* | CGC, Waterston lab |
| OS14282 | *stIs10501 [ceh-36::H1-wCherry::his-24::mCherry + unc-119(+)];wgIs707 [sptf-1::TY1::EGFP::3xFLAG + unc-119(+)]* (May still have *unc-119(tm4063) III* or *unc-119(ed3) III*) | This study, generated from strains from Waterston lab |
| OP707 | *unc-119(tm4063) III; wgIs707 [sptf-1::TY1::EGFP::3xFLAG + unc-119(+)]* | CGC, Waterston lab |
| OS1914 | *nsIs105 [hlh-17p::gfp] I* | Shaham lab(Yoshimura et al. 2008) |
| OS8753 | *nsIs105 [hlh-17p::gfp] I; eri-1 (mg366)* | Shaham lab(Yoshimura et al. 2008)  Ruvkun lab(Kennedy et al. 2004) |
| MT156 | *lin-26(n156) II* | CGC(Labouesse et al. 1996) |
| OS15214 | *nsIs105 [hlh-17p::gfp] I; lin-26(n156) II* | This study |
| VC1238 | *ztf-29(gk561) III* | CGC(The C. elegans Deletion Mutant Consortium 2012) |
| OS15122 | *nsIs105 [hlh-17p::gfp] I; ztf-29(gk561) III* | This study |
| OS15508 | *nsIs105 [hlh-17p::gfp] I; ztf-29(gk561) III; nsEx7511[pKB001(**ztf-29p(3kb)::ztf-29::ztf-29 3'UTR) + elt-2::mCherry]* | This study |
| OH7323 | *ceh-43(ot406) III; vtIs1* *[dat-1p::GFP + rol-6(su1006)] vsIs33 [dop-3::RFP] V* | CGC(Doitsidou et al. 2013) |
| OS14820 | *nsIs105 [hlh-17p::gfp] I; ceh-43(ot406) III* | This study |
| OS14781 | *nsIs105 [hlh-17p::gfp] I; ceh-43(ot406) III; nsIs975 [mir-228p::2xNLS::iBlueberry + unc-122::RFP] X* | This study |
| OS13802 | *nsIs105 [hlh-17p::gfp] I; nsIs975 [mir-228p::2xNLS::iBlueberry + unc-122::RFP] X* | This study |
| OH10279 | *ceh-43(tm480) III; otIs287 [rab-3::NLS::YFP + rol-6(su1006)]; norEx41 [ceh-43 fosmid + dat-1::mCherry]* | CGC(Doitsidou et al. 2013), Mitani lab |
| PHX5073 | *ceh-43(syb5073[ceh-43::SL2::GFP::H2B]) III* | CGC(Reilly et al. 2022) |
| OS14922 | *ceh-43(syb5073[ceh-43::SL2::GFP::H2B]) III; nsIs975 [mir-228p::2xNLS::iBlueberry) + unc-122::RFP] X* | This study |
| OS15096 | *nsIs105 [hlh-17p::gfp] I; ceh-43(ot406) III; nsEx7406[ceh-43 fosmid (WRM0633cC03) + elt-2::mCherry]* (Line 1) | This study |
| OS15097 | *nsIs105 [hlh-17p::gfp] I; ceh-43(ot406) III; nsEx7407[ceh-43 fosmid (WRM0633cC03) + elt-2::mCherry]* (Line 2) | This study |
| OS15098 | *nsIs105 [hlh-17p::gfp] I; ceh-43(ot406) III; nsEx7408[ceh-43 fosmid (WRM0633cC03) + elt-2::mCherry]* (Line 3) | This study |
| OS15331 | *nsIs105 [hlh-17p::gfp] I; ceh-43(ot406) III; nsIs975 [mir-228p::2xNLS::iBlueberry + unc-122::RFP] X; nsEx7452 [ceh-43p::ceh-43 + pSL037 (nhr-2p::his-24::mCherry) + elt-2::mCherry]* (Line 1) | This study |
| OS15332 | *nsIs105 [hlh-17p::gfp] I; ceh-43(ot406) III; nsIs975 [mir-228p::2xNLS::iBlueberry + unc-122::RFP] X; nsEx7453 [ceh-43p::ceh-43 + pSL037 (nhr-2p::his-24::mCherry) + elt-2::mCherry]* (Line 2) | This study |
| OS15333 | *nsIs105 [hlh-17p::gfp] I; ceh-43(ot406) III; nsIs975 [mir-228p::2xNLS::iBlueberry + unc-122::RFP] X; nsEx7454 [ceh-43p::ceh-43 + pSL037 (nhr-2p::his-24::mCherry) + elt-2::mCherry]* (Line 3) | This study |
| OS15476 | *nsIs105 [hlh-17p::gfp] I; ceh-43(ot406) III; nsIs975 [mir-228p::2xNLS::iBlueberry + unc-122::RFP] X; nsEx7498 [pros-1pS::ceh-43(cDNA) + elt-2::mCherry]* (Line 1) | This study |
| OS15477 | *nsIs105 [hlh-17p::gfp] I; ceh-43(ot406) III; nsIs975 [mir-228p::2xNLS::iBlueberry + unc-122::RFP] X; nsEx7499 [pros-1pS::ceh-43(cDNA) + elt-2::mCherry]* (Line 2) | This study |
| OS15478 | *nsIs105 [hlh-17p::gfp] I; ceh-43(ot406) III; nsIs975 [mir-228p::2xNLS::iBlueberry + unc-122::RFP] X; nsEx7500 [pros-1pS::ceh-43(cDNA) + elt-2::mCherry]* (Line 3) | This study |
| OS15479 | *nsIs105 [hlh-17p::gfp] I; ceh-43(ot406) III; nsIs975 [mir-228p::2xNLS::iBlueberry + unc-122::RFP] X; nsEx7501 [pSL056 (sox-2p::ceh-43 cDNA::SL2::his-24::mCherry) + pSL046 (nhr-2p::his-24:TagBFP(I174A)) + elt-2::mCherry]* (Line 1) | This study |
| OS15480 | *nsIs105 [hlh-17p::gfp] I; ceh-43(ot406) III; nsIs975 [mir-228p::2xNLS::iBlueberry + unc-122::RFP] X; nsEx7502 [pSL056 (sox-2p::ceh-43 cDNA::SL2::his-24::mCherry) + pSL046 (nhr-2p::his-24:TagBFP(I174A)) + elt-2::mCherry]* (Line 2) | This study |
| OS15481 | *nsIs105 [hlh-17p::gfp] I; ceh-43(ot406) III; nsIs975 [mir-228p::2xNLS::iBlueberry + unc-122::RFP] X; nsEx7503 [pSL056 (sox-2p::ceh-43 cDNA::SL2::his-24::mCherry) + pSL046 (nhr-2p::his-24:TagBFP(I174A)) + elt-2::mCherry]* (Line 3) | This study |
| OS15482 | *nsIs105 [hlh-17p::gfp] I; ceh-43(ot406) III; nsIs975 [mir-228p::2xNLS::iBlueberry + unc-122::RFP] X; nsEx7504 [pSL047 (mir-228p::ceh-43 cDNA::SL2::his-24::mCherry) + elt-2::mCherry]* (Line 1) | This study |
| OS15483 | *nsIs105 [hlh-17p::gfp] I; ceh-43(ot406) III; nsIs975 [mir-228p::2xNLS::iBlueberry + unc-122::RFP] X; nsEx7504 [pSL047 (mir-228p::ceh-43 cDNA::SL2::his-24::mCherry) + elt-2::mCherry]* (Line 2) | This study |
| OS15484 | *nsIs105 [hlh-17p::gfp] I; ceh-43(ot406) III; nsIs975 [mir-228p::2xNLS::iBlueberry + unc-122::RFP] X; nsEx7504 [pSL047 (mir-228p::ceh-43 cDNA::SL2::his-24::mCherry) + elt-2::mCherry]* (Line 3) | This study |
| OS15485 | *nsIs105 [hlh-17p::gfp] I; ceh-43(ot406) III; nsIs975 [mir-228p::2xNLS::iBlueberry + unc-122::RFP] X; nsEx7504 [pSL047 (mir-228p::ceh-43 cDNA::SL2::his-24::mCherry) + elt-2::mCherry]* (Line 4) | This study |
| OS15197 | *nsIs105 [hlh-17p::gfp] I; ceh-43(ot406) III; nsIs975 [mir-228p::2xNLS::iBlueberry + unc-122::RFP] X; nsEx7431 [ceh-43 fosmid (WRM0633cC03) + pSL037 (nhr-2p::his-24::mCherry) + elt-2::mCherry]* | This study |
| OS15437 | *nsIs105 [hlh-17p::gfp] I; ceh-43(tm480) III; nsIs975 [mir-228p::2xNLS::iBlueberry + unc-122::RFP] X; nsEx7450 [ceh-43 fosmid (WRM0633cC03) + pSL037 (nhr-2p::his-24::mCherry) + elt-2::mCherry]* | This study |
| OS15465 | *nsIs105 [hlh-17p::gfp] I; nsEx7490 [pSL048 (Phsp16.41::ceh-43 cDNA::SL2::his-24::mCherry::let-858 3'UTR) + elt-2::mCherry]* | This study |
| OS7698 | *nsIs408[ceh-24::GFP + odr-1::RFP]* | Shaham lab(Rapti et al. 2017) |
| OS15207 | *ceh-43(ot406) III; nsIs408 [ceh-24::GFP + odr-1::RFP]* | This study, Shaham lab(Rapti et al. 2017) |
| OH99 | *mgIs18 [ttx-3p::GFP] IV* | CGC |
| OS15298 | *mgIs18 [ttx-3p::GFP] IV; nsIs978 [mir-228p::2xNLS::iBlueberry + unc-122::RFP]* | This study |
| OS15212 | *ceh-43(ot406) III; mgIs18 [ttx-3p::GFP] IV* | This study |
| OS15299 | *ceh-43(ot406) III; mgIs18 [ttx-3p::GFP] IV; nsIs978 [mir-228p::2xNLS::iBlueberry + unc-122::RFP]* | This study |
| OS15351 | *ceh-43(ot406) III; mgIs18 [ttx-3p::GFP] IV; nsIs978 [mir-228p::2xNLS::iBlueberry + unc-122::RFP]; nsEx7459 [ceh-43p::ceh-43 + pSL037 (nhr-2p::his-24::mCherry) + elt-2::mCherry]* (Line 1) | This study |
| OS15352 | *ceh-43(ot406) III; mgIs18 [ttx-3p::GFP] IV; nsIs978 [mir-228p::2xNLS::iBlueberry + unc-122::RFP]; nsEx7460 [ceh-43p::ceh-43 + pSL037 (nhr-2p::his-24::mCherry) + elt-2::mCherry]* (Line 2) | This study |
| OS15353 | *ceh-43(ot406) III; mgIs18 [ttx-3p::GFP] IV; nsIs978 [mir-228p::2xNLS::iBlueberry + unc-122::RFP]; nsEx7461 [ceh-43p::ceh-43 + pSL037 (nhr-2p::his-24::mCherry) + elt-2::mCherry]* (Line 3) | This study |
| OS15367 | *syb9593 [hsp-16.2p::lox::degron::dam::vhhGFP4::unc-54 3'UTR; unc-119(+)] II; nsIs975 [mir-228p::2xNLS::iBlueberry + unc-122::RFP] X; nsIs1044 [hlh-17p::nls::tagrfp]* | This study (*syb9593* is modified from IS3675 from the Peter Meister lab)(Gómez-Saldivar et al. 2020) |
| OS15368 | *syb9593 [hsp-16.2p::lox::degron::dam::vhhGFP4::unc-54 3'UTR; unc-119(+)] II; ceh-43(syb9080[ceh-43::3xGAS::GFP::AID]) III; nsIs975 [mir-228p::2xNLS::iBlueberry + unc-122::RFP] X; nsIs1044 [hlh-17p::nls::tagrfp]* | This study (*syb9593* is modified from IS3675 from the Peter Meister lab)(Gómez-Saldivar et al. 2020) |
| OS15160 | *nsIs978 [mir-228p::2xNLS::iBlueberry + unc-122::RFP]; nsIs616 [glt-1p::gfp; unc-122::RFP]* | This study, Shaham lab(Katz et al. 2019) |
| OS15162 | *ceh-43(ot406) III; nsIs978 [mir-228p::2xNLS::iBlueberry + unc-122::RFP]; nsIs616 [glt-1p::gfp; unc-122::RFP]* | This study, Shaham lab(Katz et al. 2019) |
| OS14597 | *ujIs113 [pie-1p::mCherry::H2B::pie-1 3'UTR + nhr-2p::his-24::mCherry::let-858 3'UTR + unc-119(+)] II; wgIs500 [ceh-26::TY1::EGFP::3xFLAG + unc-119(+)]* | This study, generated from strains from Murray lab(Zacharias et al. 2015) and the Regulatory Element Project of modENCODE(Sarov et al. 2006; Niu et al. 2011) |
| SYS443 | *ces-2(dev112([mNeonGreen::ces-2]) I; ujIs113 [pie-1p::mCherry::H2B::pie-1 3'UTR + nhr-2p::mCherry::his-24::let-858 3’UTR + unc-119(+)] II* | CGC, Du lab(Ma et al. 2021) |
| SYS516 | *ujIs113 [pie-1p::mCherry::H2B::pie-1 3'UTR + nhr-2p::mCherry::his-24::let-858 3’UTR + unc-119(+)] II; sox-2(dev153([sox-2::mNeonGreen]) X* | CGC, Du lab(Ma et al. 2021) |
| SYS473 | *lim-7(dev125([mNeonGreen::lim-7]) I; ujIs113 [pie-1p::mCherry::H2B::pie-1 3'UTR + nhr-2p::mCherry::his-24::let-858 3’UTR + unc-119(+)] II* | CGC, Du lab(Ma et al. 2021) |
| CA1200 | *ieSi57 [eft-3p::TIR1::mRuby::unc-54 3'UTR + Cbr-unc-119(+)] II; unc-119(ed3) III* | CGC, Dernburg lab(Zhang et al. 2015) |
| OS15343 | *nsIs105 [hlh-17p::gfp] I; ieSi57 [eft-3p::TIR1::mRuby::unc-54 3'UTR + Cbr-unc-119(+)] II; ceh-43(syb9080[ceh-43::3xGAS::GFP::AID]) III* | This study |
| OS15344 | *nsIs105 [hlh-17p::gfp] I; ieSi57 [eft-3p::TIR1::mRuby::unc-54 3'UTR + Cbr-unc-119(+)] II* | This study |
| OS15345 | *nsIs105 [hlh-17p::gfp] I; ceh-43(syb9080[ceh-43::3xGAS::GFP::AID]) III* | This study |
| OS15346 | *ieSi57 [eft-3p::TIR1::mRuby::unc-54 3'UTR + Cbr-unc-119(+)] II; ceh-43(syb9080[ceh-43::3xGAS::GFP::AID]) III* | This study |
| OS15505 | *ieSi57 [eft-3p::TIR1::mRuby::unc-54 3'UTR + Cbr-unc-119(+)] II; ceh-43(syb9080[ceh-43::3xGAS::GFP::AID]) III; nsIs978 [mir-228p::2xNLS::iBlueberry + unc-122::RFP]; nsIs616 [glt-1p::gfp; unc-122::RFP]* | This study |
| OS15503 | *ieSi57 [eft-3p::TIR1::mRuby::unc-54 3'UTR + Cbr-unc-119(+)] II; nsIs978 [mir-228p::2xNLS::iBlueberry + unc-122::RFP]; nsIs616 [glt-1p::gfp; unc-122::RFP]* | This study |
| OS15504 | *ceh-43(syb9080[ceh-43::3xGAS::GFP::AID]) III; nsIs978 [mir-228p::2xNLS::iBlueberry + unc-122::RFP]; nsIs616 [glt-1p::gfp; unc-122::RFP]* | This study |
| OS15502 | *ieSi57 [eft-3p::TIR1::mRuby::unc-54 3'UTR + Cbr-unc-119(+)] II; ceh-43(syb9080[ceh-43::3xGAS::GFP::AID]) III; nsIs978 [mir-228p::2xNLS::iBlueberry + unc-122::RFP]* | This study |
| OS15540 | *glt-1(ns1117[glt-1::SL2::GFP::H2B]) X* | This study |
| OS15542 | *glt-1(ns1119ns1117[glt-1::SL2::GFP::H2B]) X* | This study |
